# Analytical characterization of Parylene-C degradation mechanisms on Utah arrays: evaluation of in vitro Reactive Accelerated Aging model compared to multiyear *in vivo* implantation

**DOI:** 10.1101/743831

**Authors:** Ryan Caldwell, Matthew G. Street, Rohit Sharma, Pavel Takmakov, Brian Baker, Loren Rieth

## Abstract

Implantable neural microelectrodes are integral components of neuroprosthetic technologies and can transform treatments for many neural-mediated disorders. However, dielectric material degradation during long-term (> 1 year) indwelling periods restricts device functional lifetimes to a few years. This comprehensive work carefully investigates *in vivo* material degradation and also explores the ability of *in vitro* Reactive Accelerated Aging (RAA) to evaluate implant stability. Parylene C-coated Utah electrode arrays (UEAs) implanted in feline peripheral nerve for 3.25 years were explanted and compared to RAA-processed devices, aged in phosphate buffered saline (PBS) + 20 mM H_2_O_2_ at either 67 or 87 °C (28 or 7 days, respectively). Electron microscopy revealed similar physical damage characteristics between explants and RAA (87° C) devices. Parylene C degradation was overwhelmingly apparent for UEAs from both RAA cohorts. Controls aged in PBS alone displayed almost no damage. Spectroscopic characterization (EDX, XPS, FTIR) found clear indications of oxidation and chlorine abstraction for parylene C aged *in vivo*. While *in vitro* aging was also accompanied by signs of oxidation, changes in the chemistry *in vivo* and *in vitro* were statistically different. Analysis of RAA- aged devices identified UEA fabrication approaches that may greatly improve device resistance to degradation. This work underscores the need for an improved understanding of *in vivo* damage mechanisms, to facilitate the critical need for representative *in vitro* accelerated testing paradigms for long-term implants.

## **1.** Introduction

The performance of medical devices is linked to the materials used in their fabrication, with material degradation resulting in device failure. Material degradation has been identified as one of the key factors limiting the lifetime of neural electrodes such as the Utah array, and the mechanism for this degradation have not been thoroughly characterized using analytical techniques.. The development of robust electrode technologies is an active research pursuit, as these devices have the potential to improve the lives of millions living with disabilities such as limb loss, spinal cord injury, and motor deficits [1]– [4]. However, implantable neural microelectrodes have yet to reliably demonstrate continuous functionality for 10 years or more *in vivo*, the accepted metric for clinical viability [5]. Meeting this metric will require improved materials and fabrication methods informed by rigorous (accelerated) testing of architectures and materials. The current paradigm for neural implant development focuses on collecting medical device performance data using *in vivo* testing in large animals as model organisms [6]. While, this approach is an important pre-clinical step for evaluating device safety and efficacy, it is impractical for rapid and high-throughput testing of material reliablity during early development stages [7].

A common testing paradigm for quick assessment of implantable device reliability is *in vitro* soak testing in saline solutions. This test is often performed at temperatures of 37°C or higher to represent exposure *in vivo* in real-time or accelerated aging, respectively [8]. Such testing evaluates water ingress, dissolution/hydrolysis reactions, corrosion, hydration/swelling, and changes in material properties that occur in the *in vivo* environment. These tests have been a useful first step in determining the robustness of exposed materials as they are relatively easy to implement and interpret. The material performance of neural microelectrodes has been commonly assessed in this manner [9]–[15]. Furthermore, the lifetime of encapsulation strategies against fluid ingress and degradation can be assessed through impedance measurements to test improved encapsulation materials and processes, and inform better neural electrode design.

However, *in vitro* saline testing *does not* represent important degradation mechanisms present *in vivo* due to the local millieu having many components beyond saline. Therefore, *in vitro* testing does not represent important degradation mechanisms observed from long-term implants, hampering the utility of these tests to drive material optimization research and quantify device lifetime. For example, Hämmerle *et al.* found that a thermally grown silicon dioxide dielectric layer that remained unchanged after 21 months *in vitro* nevertheless completely etched away *in vivo* after 10 months, leading to failure of an implanted photodiode retinal prosthetic [12]. Polyimide-insulated tungsten microwire arrays are known to degrade *in vivo* [16], but this degradation is not predicted by aging in physiological saline alone [17].

Parylene C (PPX-C) is an important biopolymer for medical devices due to its conformal deposition, low dielectric constant and moisture permeation, and USP Class VI classification [11], [15], [18]–[22]. Numerous reports of testing PPX-C lifetime *in vitro* [14], [19], [23] have failed to replicate the damage mechanisms reported from *in vivo* studies, including cracking, delamination, erosion, and cratering observed by our lab and others [24]–[27]. The underlying causes of such degradation need to be elucidated, and novel *in vitro* test beds that represent them are needed verify mechanisms and enable better accelerated aging tools.

This study investigates the addition of oxidative species in saline test to represent these degradation mechanisms using advanced analytical techniques. Takmakov *et al*. have developed and published on an oxidative reactive accelerated aging system (RAA) for accelerated aging of neural microelectrodes, by soaking in an 87°C bath composed of PBS with 20 mM H_2_O_2_ [17]. Oxidation species are known to be generated by activated microglia, macrophages, and neutrophils engaged as part of the foreign body response [28]. When activated, these cells release reactive oxygen species (ROS) such as hydrogen peroxide (H_2_O_2_), superoxide anion (•O_2_^-^), and hydroxyl radical (•OH). Signs of oxidative stress have been observed to persist at neural implant sites for >8-16 weeks [29], [30], and long-term exposure to such conditions may degrade microelectrode materials.

*In vitro* studies incorporating H_2_O_2_, the most readily replicated and controlled ROS [31], have noted accelerated tungsten microwire electrode corrosion [17], [31], similar to corrosion observed *in vivo* [16]. While Takmakov *et al.* observed impedance decreases for all four microelectrode types tested, as well as damage to polyimide, little physical PPX-C degradation was initially observed by electron microscopy. This contrasts with prior reports of *in vivo* PPX-C damage [17]. Parylene polymers undergo thermal and photolytic oxidation [32], [33], and parylene damage from thermal oxidation has been proposed to occur via the transformation of methylene bonds to ester bonds, [34]. Oxidation-induced ester groups in PPX-C would be vulnerable to hydrolysis *in vivo* [35]–[37], and may be a mechanism that contributes to PPX-C damage. A closer evaluation of oxidation effects from PPX-C aging is warranted.

This study extends RAA testing to evaluate kinetics of PPX-C degradation through testing at both 87 °C and 67 °C, and establishes mechanisms of degradation through detailed characterization of a large sample size of devices (total of 43). We have found that PPX-C degradation in RAA testing does indeed repeatably occur, and exhibits similarities to *in vivo* degradation. Furthermore, these results are compared to a unique new analytical data from two Utah slanted electrode arrays (USEAs) explanted after 3.25 years’ dwell time in feline femoral and sciatic nerve [38]. These devices exhibit PPX-C damage characteristics similar to those of previous reports, suggesting that insights into degradation mechanisms can be determined from analytical characterization data.

A key aspect of our study is the complementary set of analytical characterization techniques used to probe the chemistry of damaged PPX-C films. Secondary electron microscopy with energy dispersive X-ray analysis (SEM/EDX) and electrochemical measurements, *e.g.* impedance (for example, see [16], [17], [24], [27], [39], [40]) have been used to gain important insights into degradation process. However, they provide limited information regarding degradation chemistry. Spectroscopic techniques can probe film chemistry, but have not been used on neural microelectrode thin films, partly due to difficult sample preparation [15]. This report uses novel methods to prepare and characterize samples using spectroscopic analysis such as energy dispersive X-ray spectroscopy (EDS), X-ray photoelectron spectroscopy (XPS), and Fourier transform infrared spectroscopy (FTIR) of PPX-C films on UEAs and USEAs. Our results suggest that oxidation of PPX-C films occurs *in vivo*, and that *in vivo* and *in vitro* film damage may be influenced by device fabrication protocols that can facilitate oxidation. This work enhances the understanding of materials degradation that has been reported in previous studies, and will help inform the design and testing of neural microelectrodes and other implantables to improve their long-term performance reliability.

## 2. Materials and methods

Devices characterized in this study included 2 USEAs explanted after 3.25 years in feline peripheral (sciatic and femoral) nerve, as well as 20 UEAs aged using the RAA-protocol. An additional 8 control UEAs were aged in PBS (without H_2_O_2_), and test structures including interdigitated electrodes (IDEs) and test UEAs (t-UEAs) were aged in RAA and control conditions to provide supplementary data. Table 1 summarizes the experimental protocol. The USEAs were part of a separate study into functional nerve stimulation [38], so pre-implant characterization data are limited. All UEAs were characterized before *in vitro* aging (RAA) using scanning electron microscopy (SEM) and electrochemical impedance spectroscopy (EIS). All USEAs and UEAs were characterized post-aging using SEM, EIS, EDS, XPS, and FT-IR, as described below. To comprehensively characterize the degradation mechanisms, 7 different types of reference samples were required for baseline and to control for sample preparation methods. Fig. 1 shows an illustration of a UEA, as well as the novel sample preparation techniques that were developed.

**Fig. 1.**
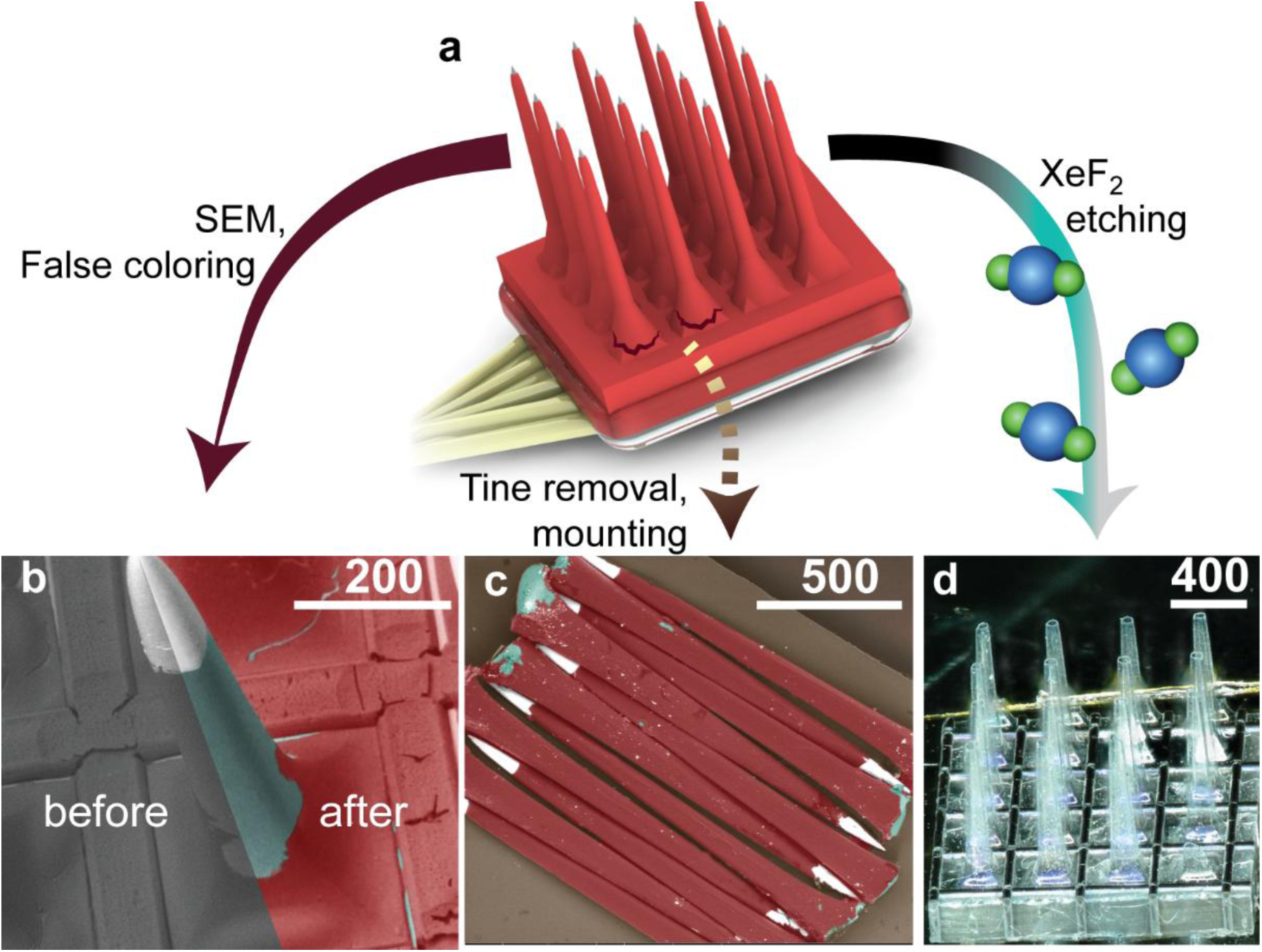
Illustrations and micrographs of UEAs and sample processing methods. (a) Diagram of UEA, highlighting processing techniques shown in (b)-(d). (b) Micrograph of an electrode tine demonstrates false coloring to highlight iridium oxide (white), silicon (blue-green), and PPX-C (red) based on extensive experience regarding contrast mechanisms and EDS chemical analysis from raw images (“before”) to false colored (“after”). (c) Electrode tines were detached and laid tip-to-base to form a pseudo-planar surface for spectroscopy. (d) Optical micrograph of hollow PPX-C sheaths from UEA after silicon removal via XeF_2_ etching, placed on adhesive like (c) for FTIR analysis.

**Table 1.**
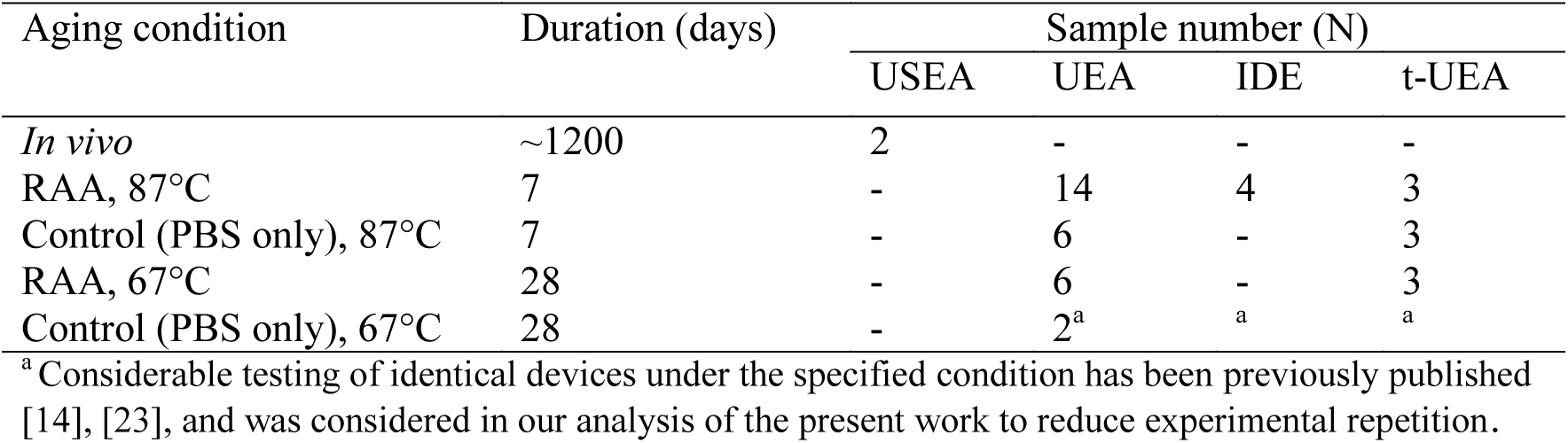
Summary of RAA experimental design.

| Aging condition | Duration (days) | Sample number (N) |  |  |  |
| --- | --- | --- | --- | --- | --- |
|  |  | USEA | UEA | IDE | t-UEA |
| *In vivo* | ~1200 | 2 | - | - | - |
| RAA, 87°C | 7 | - | 14 | 4 | 3 |
| Control (PBS only), 87°C | 7 | - | 6 | - | 3 |
| RAA, 67°C | 28 | - | 6 | - | 3 |
| Control (PBS only), 67°C | 28 | - | 2<sup>a</sup> | <sup>a</sup> | <sup>a</sup> |

<sup>a</sup> Considerable testing of identical devices under the specified condition has been previously published [14], [23], and was considered in our analysis of the present work to reduce experimental repetition.

### 2.1 UEA fabrication and PPX-C encapsulation

UEA construction has been reported at length [41], [42] and proceeded from *p*-type silicon (0.01- 0.05 Ω cm, Virginia Semiconductor, Fredericksburg, VA), which was diced, processed, and etched to create 4×4 arrays of sharp silicon tines 1 or 1.5 mm in length, electrically isolated by glass-filled kerfs. Iridium oxide was deposited on tips of the silicon tines by reactive sputter deposition [43], [44]. Platinum was sputter deposited and lithographically patterned on the planar UEA backside to create wire bond pads.

After metal deposition, devices were encapsulated with PPX-C. The encapsulation process began with 2 hours’ exposure to vapor-phase A-174 silane adhesion promoter (Momentive Performance Materials, Waterford, NY). PPX-C films of 5-6 µm thickness were then deposited from parylene C dimer (DPX-C) in a PDS 2010 (both from Specialty Coating Systems, Indianapolis, IN). Film thickness was verified on witness substrates by scoring the film and measuring step height with a profilometer (Tencor, Milpitas, CA). Adhesion tests were performed on silicon monitor wafer pieces using the ASTM D3359 protocol (tape test), and acceptable scores of 4B-5B [45] were found for all deposition runs.

UEA electrode tips were exposed by etching PPX-C using an oxygen plasma deinsulation process. The UEAs were packaged in a second aluminum foil mask such that 50-100 µm of the tips penetrated through the foil, and etched in O_2_ plasma (March Plasma Systems, Inc., Concord, CA) until the exposed PPX-C was removed. The etch durations were typically 18-24 minutes at 100 W and 0.4 Torr, and parylene removal was verified through backscatter electron microscopy (BSEM) owing to the high Z-contrast between iridium oxide and parylene.

After deinsulation, UEAs were manually wire bonded (West Bond, Anaheim, CA) to gold- flashed printed circuit boards (Circuit Graphics, Salt Lake City, UT) using polyester enamel-insulated gold bonding wire (99% Au:1% Pd) (Sandvik, Stockholm, Sweden). Bond pads and wires were overmolded with MED-4211 silicone (NuSil, Carpenteria, CA) for electrical insulation and physical reinforcement.

### 2.2 Test structure fabrication

Test structures permitted additional characterization of PPX-C and encapsulation damage from *in vitro* aging. Two types of structures were used: interdigitated electrodes (IDEs, N=4 total) and test arrays (t-UEAs, N=9 total). IDEs were fabricated from fused silica wafers (Hoya, Tokyo, Japan) and encapsulated with PPX-C, as previously described [14], [23], [13]. The simplicity of IDE test structures greatly reduced the degree of sample preparation needed for spectroscopy, and planar surfaces provided for a sensitive measure of changes to encapsulation impedance. The t-UEAs were fabricated from unmetallized UEAs, which were mounted to insulated copper wire using silicone and completely encapsulated with PPX-C. Specifically, the tips of t-UEAs were not exposed through PPX-C through oxygen plasma processing. Therefore, t-UEAs differed from UEAs in that they were not subject to metal deposition, oxygen plasma, or wire bonding. Comparison of PPX-C aging on t-UEAs with that of fully fabricated UEAs enabled detection of manufacturing-induced effects, particularly the impact of oxygen plasma processing. IDEs were not ideal for this purpose as they lacked UEA topography, which we have shown to be an important factor to consider when evaluating encapsulation performance [14].

### 2.3 Reactive accelerated aging (RAA)

The materials and methods for RAA aging have been carefully reported [17], [46] and occurred in a jacketed flask (Pine Research Instrumentation, Durham, NC) filled with 20 mM H_2_O_2_ in PBS, prepared using deionized water and prefabricated PBS tablets (Sigma Aldrich, St. Louis, MO). Solution temperature was controlled using a Haake C-25 recirculating heater with mineral oil (Fisher Scientific, Pittsburgh, PA). Thermostat set points were slightly above the target to achieve flask conditions that did not vary more than 1°C from desired temperatures, verified with a glass thermometer. RAA was performed at 67 °C and 87 °C, corresponding to respective nominal PBS soak testing (non-RAA) aging acceleration factors *f* of 8× and 32× at those temperatures [14], [23], [13], compared to physiological conditions. It is important to note that these acceleration factors are derived from empirical Arrhenius modeling of accelerated reactions [8]. The temperature of 87° was chosen as the highest acceptable accelerating aging temperature, based on prior work [17]. Aging at 67 °C was performed to investigate the underlying reaction kinetics for the degradation of PPX-C, through comparisons between results obtained at the two temperatures. A closed loop control system with continuous direct UV measurement was developed to maintain H_2_O_2_ levels during the lengthier 67 °C run as previously described [47] (for new closed loop RAA with electrochemical feedback see [48]). Extensive aging of both UEAs and IDEs in PBS at 67 °C has been previously reported and provided as a dataset for comparison to the results of this study [14].

All devices which underwent *in vitro* aging were fixed to custom-machined 24/40 PTFE flask stoppers [47] with silicone, to allow their placement in the RAA solution and prevent evaporation. Up to four stoppers and their associated samples were placed in the RAA system simultaneously to maximize uniformity of exposure conditions. Control devices were aged in PBS alone at 87°C and 67°C for minimum durations of 7 and 28 days, respectively, to elucidate the effects of ROS relative to saline soaking. This also helped determine if the higher 87 °C aging temperature activated additional failure mechanism through exceeding the PPX-C glass transition temperature (T_g_, 35-150°C [49]–[51]).

### 2.4 USEA explant details

The explanted USEAs described in this study were used to record and stimulate motor and sensory neural activity. Details regarding the objective, surgical procedures, and experimental protocols for the implants are reported elsewhere [38]. Briefly, three 10×10 USEAs were fabricated with IrO_x_ tip metal and PPX-C of 2.8 µm nominal thickness. Ethylene oxide (EtO) sterilization occurred at University of Utah hospital. Devices were implanted in the left sciatic and femoral nerves of an adult female cat under protocols approved by the University of Utah Institutional Animal Care and Use Committee. After 3.25 years, the feline was euthanized with intravenous saturated KCl, and two arrays were carefully extracted from unfixed tissue. To remove tissue residue, USEAs were soaked overnight in ENZOL® enzymatic detergent prepared according to manufacturers’ instructions (Johnson & Johnson, New Brunswick, NJ), followed by a gentle rinse in deionized water.

### 2.5 Electrochemical impedance spectroscopy (EIS)

UEAs were electrochemically characterized via two-electrode EIS utilizing a large platinum wire (surface area > 24 mm^2^) as both counter and reference. Measurements were conducted with a Reference 600 potentiostat (Gamry Instruments, Warminster, PA) from 1-10^5^ Hz at 10 points per decade, utilizing a 25 mV RMS excitation signal. PBS for electrochemical measurements was prepared with 140 mM NaCl, 2.6 mM KCl, 1.8 mM KH_2_PO_4_, and 10 mM Na_2_HPO_4_.

### 2.6 Electron microscopy and energy dispersive X-ray spectroscopy (EDS)

All USEAs, UEAs, and most test structures were imaged with a Quanta 600 FEG using secondary electron (SE) and backscatter electron (BSE) detectors in low vacuum mode (FEI, Hillsboro, OR). Samples were imaged at 0.15 Torr with 15 kV primary beam energy, and no gold coating. Secondary electron imaging helped to distinguish surface topography and contamination, and backscattered electron imaging distinguished materials through atomic number (Z) contrast.

High-resolution imaging and cross section preparation for PPX-C thickness measurements was done using a Helios NanoLab 650 (Thermo) equipped with a magnetic immersion lens, SE detector, and gallium focused ion beam (FIB) column. Samples analyzed in this way included USEA explants and two UEAs aged with RAA at 87 °C. Immersion imaging parameters were a 2 kV primary beam energy, and 50 pA current. Prior to ion milling, a thin platinum strap was selectively deposited over the viewing region to reduce charging and image drift, and ∼1 µm thick platinum strap was created over the cross section location to minimize artifacts from milling. Milling occurred at 30 kV primary beam energy for the Ga source, with currents of 2.1 nA for rapid removal and 80 pA for polishing. Micrographs were enhanced in Photoshop (Adobe, San Jose, CA) to highlight materials identified from combined information from secondary electron and backscatter micrographs, on-site spectroscopic measurements (EDS analysis in particular), and extensive experience with UEA imaging. Delineations between iridium oxide, silicon, PPX-C, silicone, and other materials were highlighted with a consistent false color scheme, as shown in Fig. 1b. Raw and uncolored images are provided in supplementary data. EDS is a semi-quantitative technique to measure sample composition. Spectra were collected using a detector from EDAX Inc., (Mawah, NJ) on the Quanta 600 FEG for as-received samples.

Standardless spectra over 60×60 µm^2^ areas of PPX-C were collected for 50 seconds using 20 kV, spot size of 4, and 3000× magnification. Oxygen and chlorine atomic percent (at%) were normalized to carbon at% and analyzed for trends.

### 2.8 X-ray photoelectron spectroscopy (XPS)

XPS complemented EDS composition data, and provided more surface sensitive data and chemical bonding information. A major challenge was acquiring sufficient signal from UEAs, which lead to developing a sample preparation technique to approximated planar surfaces (Fig. 1c). XPS was performed over sample centers to maximize signal from parylene-coated regions. The adhesive contained oxygen complexes, requiring care in preparation, controls, references, and data analysis to prevent confounds from oxygen artifacts. Electrodes were only used in sample preparation if they exhibited >50% PPX-C coverage.

XPS spectra were collected with AXIS Ultra DLD system (Kratos Analytical, Manchester, UK). Samples were pumped overnight to ∼10^-8^ Torr prior to being introduced to analysis chamber. Argon beam sputtering for 0 (as-received), 30, and 180 seconds prior to spectra collection were used to collect spectra from the surface, after removing surface contaminants, and from within the film. Spectra were collected using a monochromatic Al K_α_ source (1486.7 eV). Survey scans were collected at a pass energy of 160 eV and 1 eV step size while high-resolution region scans were collected at 40 eV pass energy and 0.1 eV step size. Collected spectra were corrected for Shirley background signal, analyzed, and deconvoluted using CasaXPS software. High-resolution photoelectron peaks were corrected for binding energy shift by referencing the C 1s peak of aliphatic C at 284.5 eV [52]. Deconvolution of C to O bindings in the C envelop was performed by first quantifying the C to Cl ratio using the Cl envelope. This was assigned to the C envelope along with fitted aliphatic C peak, and the residual envelope area was fitted to approximations of C to O binding peaks, as well as aromatic C satellite peaks.

### 2.9 Fourier transform infrared spectroscopy (FTIR)

Attenuated total reflectance (ATR) FTIR provided sensitive detection of chemical bonds such as carbon-oxygen complexes. ATR requires clamping the PPX-C against the crystal, which was impossible while the parylene remained on UEA electrodes. Therefore, the silicon was selectively removed while preserving PPX-C film using dry isotropic XeF_2_ etching in a Xetch system (Xactix Inc., Pittsburgh, PA), at 3 Torr XeF_2_, and 4 Torr N_2_, with 18 sec dwell time and a minimum of 200 etch cycles. This preparation was destructive (Fig. 1d), so was the final method used, and created C─F complexes that had to be accounted for in FTIR measurements. The resulting PPX-C was laid on vacuum grade adhesive to form a continuous layer. Silicone-based adhesive was used for its flat absorbance spectra at 1500-2000 cm^-1^. As ATR-FTIR probe depth of ∼2 µm at 1000 cm^-1^ could exceed the thickness of damaged parylene, use of silicone adhesive ensured the integrity of carbonyl peaks near 1700 cm^-1^.

PPX-C absorbance was measured with a Nicolet 6700 (Thermo Fisher Scientific, Waltham, MA) equipped with a single bounce diamond Smart iTR accessory, a deuterated triglycine sulfate detector, and a KBr/Ge beamsplitter. Spectra were taken from 650 to 4000 cm^-1^ at 4 cm^-1^ resolution, averaged over 256 scans, and corrected for ATR spectral aberrations utilizing included functionality within OMNIC software (Thermo). Spectra were area-normalized to the C═C─C stretch peak near 1606 cm^-1^.

### 2.10 Reference samples for spectral characterization

Reference samples were used to control for and validate sample preparation methods, and aid interpretation of spectral results concerning the chemical nature of PPX-C damage. Samples typically consisted of PPX-C deposited on polished silicon or UEA substrates, according to the PPX-C deposition process already described. Additional processing is outlined in Table 2, along with the intended case each sample was meant to reference.

**Table 2.**
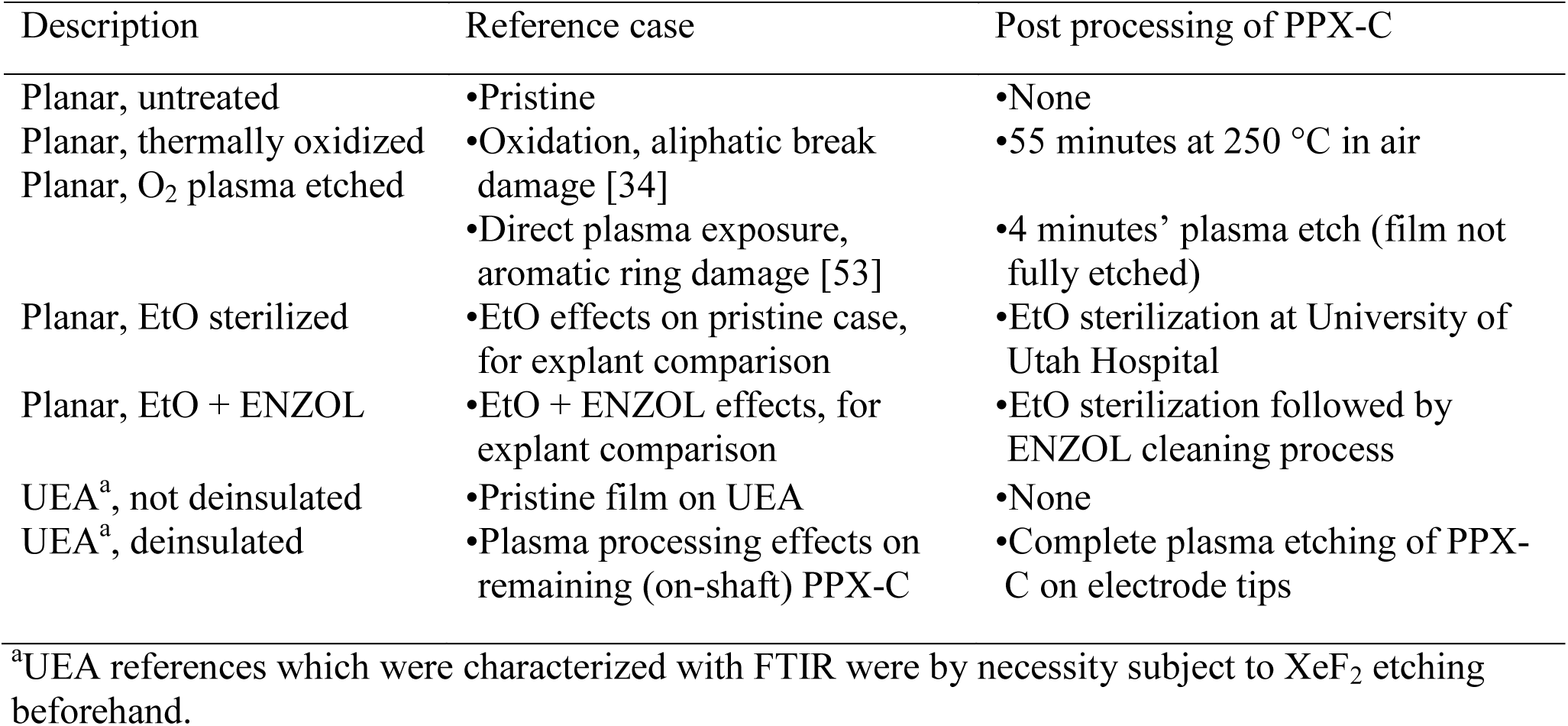
Reference samples employed for EDS, XPS, and FTIR spectroscopic characterization.

### 2.11 Statistical processing

Post-hoc statistical tests were performed on data to quantify trends in the data. The sample size of UEAs explanted after long indwelling periods was particularly limited, therefore statistical tests were not used to draw conclusions regarding the general nature of *in vivo* aging compared to RAA or other conditions. Rather, tests were only performed to strengthen our understanding of the data presented here, to inform future studies and development as well as drive improved study and test design. Analysis was performed using IBM SPSS Statistics version 24 (IBM, North Castle, NY). Groups selected for analysis were tested for significance with Welch’s ANOVA to account for unbalanced sample groups, as well as unequal variance, determined through Levene’s test. Games-Howell post-hoc tests were done to compare group means. A value of p=0.05 was considered significant.

## 3. Results

Strong similarities in surface morphology were observed via SEM between *in vivo* and 87 °C RAA UEAs. Chemical analyses revealed changes that followed clear trends, including an elevated oxygen concentration of *in vivo* samples and of UEAs aged *in vitro*, compared to untreated films. While all samples were characterized with SEM and EIS, only selected samples were subject to chemical analysis. Sample counts used for each method are given in supplemental Table S1. For brevity, detailed ethylene oxide sterilization (EtO) and EtO+ENZOL (a common enzymatic cleaner) reference sample results are omitted from this report. No indication was found using EDS, XPS, and FTIR that such processing altered film properties versus untreated references, in agreement with previous work [54]. Since the ideal accelerated aging test system will mimic material damage mechanisms observed *in vivo*, explant results were considered a benchmark for comparison of *in vitro* accelerated aging methods. We regarded aging controls as representative of the current prevailing bench top aging techniques which utilize only temperature-controlled PBS or similar electrolyte. RAA results showed how modifying such existing test methods to include oxidative chemistry drives outcomes towards *in vivo* results. As changes to PPX-C condition were the primary focus of this work, post-aging electrode metal and silicon structure are only described in brief.

### 3.1 Electron microscopy

In general, damage to arrays as observed through electron microscopy was consistent between electrodes within each array. Therefore, images of electrodes are representative of the arrays from which they originate. Sciatic and femoral explants are shown in Figs. 2 and 3, respectively (for uncolored images see Figs. S1 and S2). Differences between electrode shapes (*e.g.* tip radius) on explanted USEAs were typical for as-fabricated devices.

**Fig. 2.**
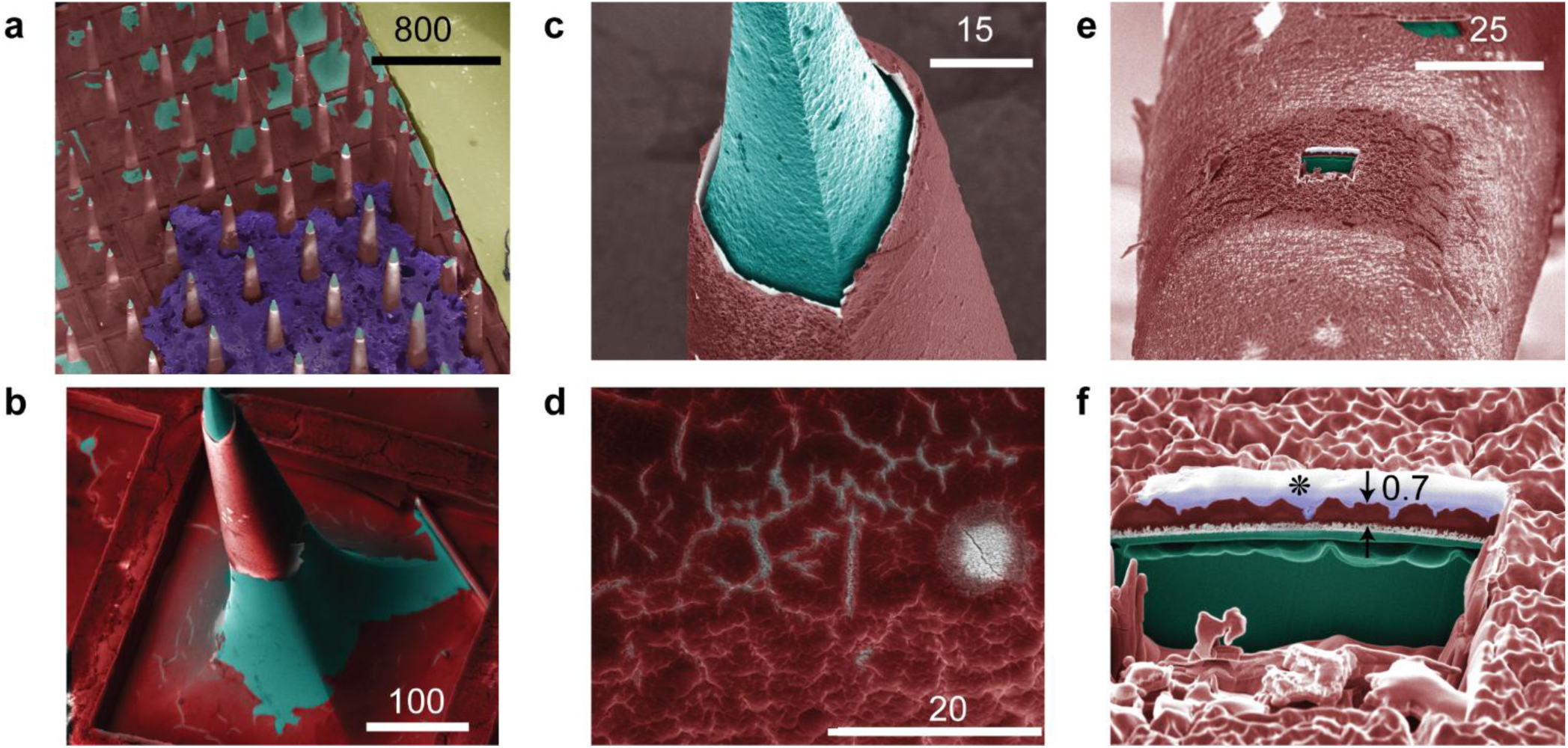
USEA explanted from cat sciatic nerve. Scale bars are in micrometers. (a) Array view showing widespread tip metal damage and PPX-C degradation. Residual tissue shown in purple, and silicone potting of gold wires shown in yellow. (b) View of single electrode, with tip metal removed and degraded PPX-C. (c) Electrode tip with silicon pitting and undercut IrO_x_/PPX-C film. (d) Detail of PPX-C over metal shows cracking and cratering. (e) Region of PPX-C along shaft that was subject to FIB cross-sectioning. (f) Detail of (e) shows considerable PPX-C surface topography, silicon pitting/etching underneath IrO_x_, and evidence of thinned PPX-C with respect to 3 µm initial thickness. In (f), light blue (marked with *) on top of parylene (red) is protective Pt deposited *in situ*. Blue-green – silicon; light gray – IrO_x_; yellow – silicone.

**Fig. 3.**
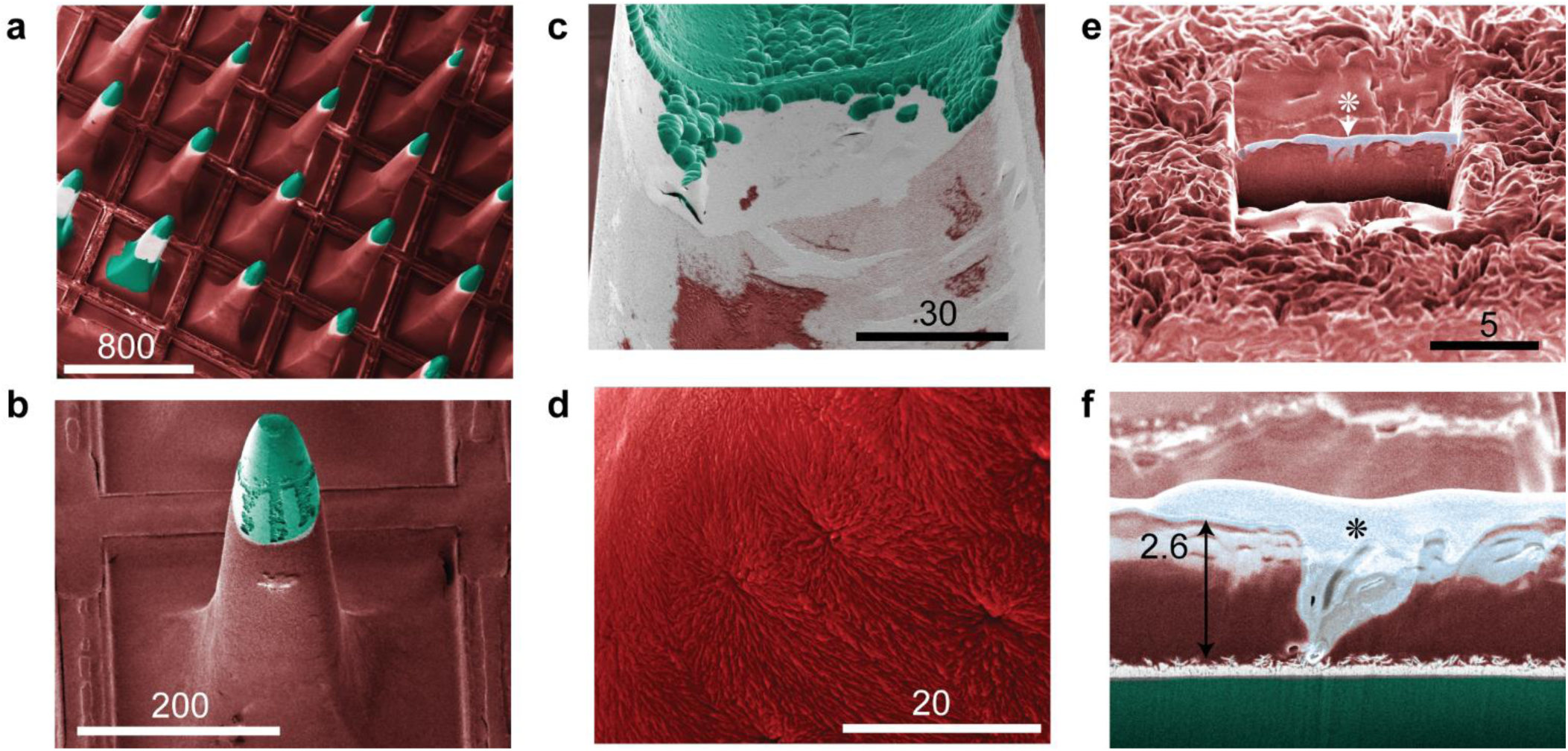
USEA explanted from cat femoral nerve. Scale bars are in micrometers. (a) Array view shows widespread electrode tip damage. PPX-C on corner electrodes was intentionally removed so that they might act as on-array references. (b) An individual electrode shows etching of silicon exposed around IrO_x_. (c) Detail of electrode tip (different from (b)) shows silicon pitting and residual PPX-C, presumably after *in vivo* degradation. (d) PPX-C over many electrodes showed surface degradation, but not complete erosion as seen in (c). (e) Considerable PPX-C surface topography visible around FIB cross section of parylene film over IrO_x_ on electrode shaft. (f) Full detail of cross section, showing PPX-C thickness varying from more than 2.6 µm to complete removal. For (e) and (f), light blue (marked with *) on top of parylene (red) is protective Pt deposited *in situ*. Blue-green – silicon; light gray – IrO_x_.

USEAs explanted from sciatic and femoral nerves showed removal of the IrO_x_ electrode metallization not protected by PPX-C, for all 200 electrodes imaged. Pitting and etching/dissolution of silicon at exposed tips was apparent for both devices as well, in many cases undercutting IrO_x_ that remained on electrode shafts underneath residual PPX-C. Cross-sectional imaging with FIB through tip metal observed the dendritic IrO_x_ structure expected of our reactive IrO_x_ sputtering process, suggesting little physical damage or change to metal that remained on electrodes. The sciatic explant cross section showed silicon pitting underneath iridium oxide film covered by PPX-C (Fig. 2f) significantly down the shank from the exposed electrode. The same was not seen for the femoral explant (Fig. 3f), from a small population of electrodes sampled.

Although similar observations of tip metal and silicon were made between USEAs, distinct PPX- C damage modes were seen for the two arrays. Both devices exhibited roughened PPX-C with fissures that completely penetrated the film, but the femoral explant in general displayed continuous PPX-C coverage of electrodes across the entire array. PPX-C thickness for this device measured via FIB cross section showed that evidence of the original 2.8 µm thick film remained, despite film roughening and fissure formation. In contrast, the sciatic explant showed exposure of silicon along the shaft or at the base of electrodes due to complete removal of PPX-C. Interestingly, residual PPX-C for most sciatic explant electrodes was present on electrode metal that was not originally exposed during oxygen plasma processing, although close observation of this film revealed cracks and craters that traversed the PPX-C film (Fig. 2d), and can therefore be expected to generate additional exposed surface and current paths. Backscatter images showed reduced contrast between residual tip metal and PPX-C appearance on the sciatic explant, compared to PPX-C and tip metal on the femoral explant, indicating a thinned PPX-C film on the former. This was confirmed with FIB cross sectional measurements, which showed PPX-C thickness on the sciatic explant being less than 1 µm compared to the original 2.8 µm thick film.

*In vitro* aged control UEAs generally exhibited no change to PPX-C encapsulation after aging in PBS, compared to before aging. A representative control array from the soak testing is shown in Fig. 4a, where the PPX-C film is observed to be completely intact. One control array out of six processed at 87 °C did exhibit nearly imperceptible PPX-C cracking, which became more visible with charging from the primary; an image of this outlier is shown in Fig. 4b. This was our first observation of PPX-C crack formation after aging in PBS, and the temperature at which it occurred (87 °C) is higher temperature than prior aging protocols (25 to 67 °C). Prior studies of UEAs aged at 67 °C in PBS by our group, both published [14] and unpublished (>10 arrays total), have never recorded PPX-C crack formation solely due to aging in PBS. However, the cracking seen here would have been difficult to detect without deliberate sample charging, a practice which was avoided during routine inspections. Although unlikely, it is possible that previous samples were incorrectly identified as damage-free. However, the samples in this report were inspected in detail, and our assessment of damage is made with confidence.

**Fig. 4.**
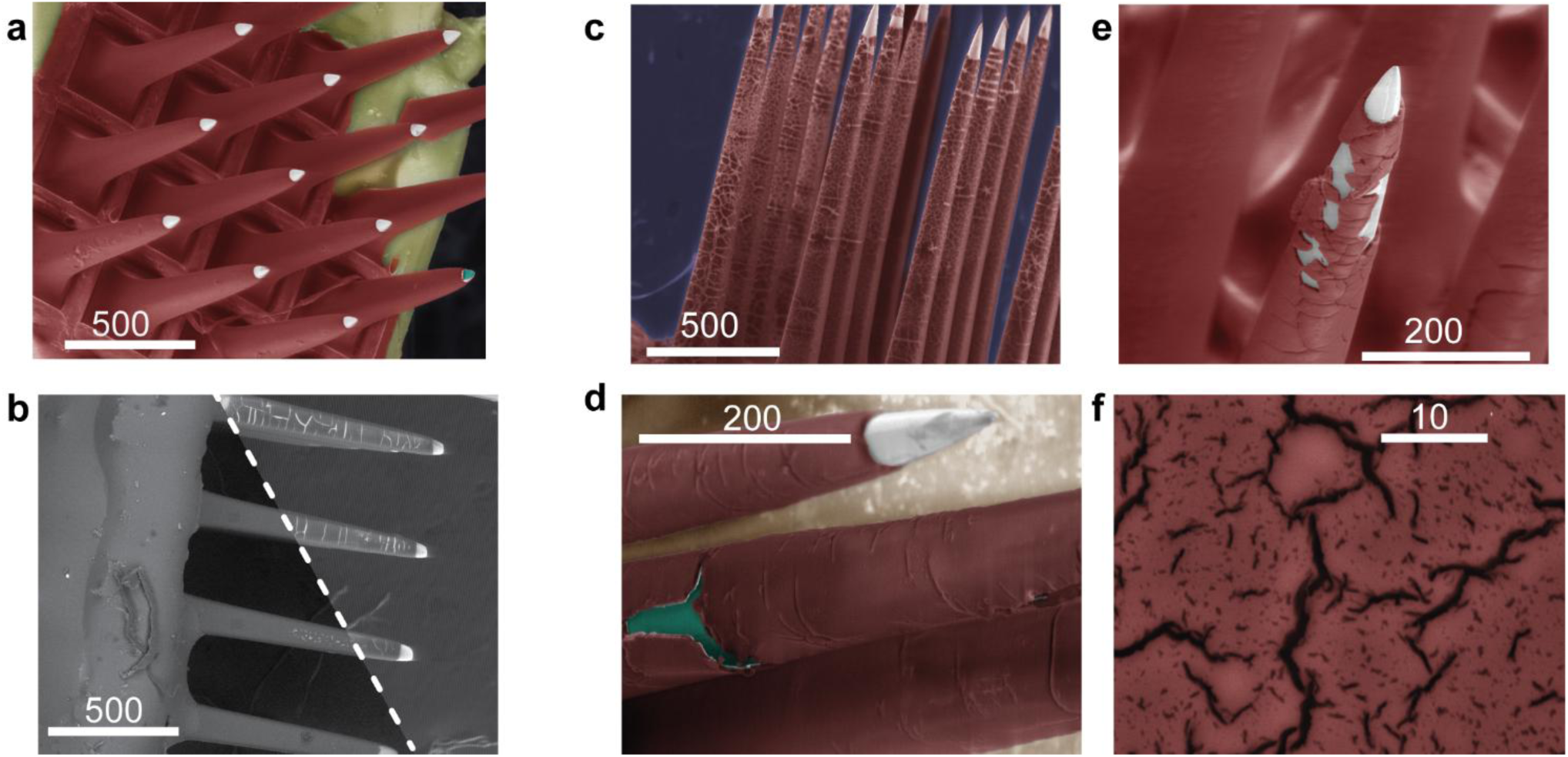
**S**EM images of control UEAs subject aged in PBS alone (a-b), and devices aged using RAA at 67 °C (c-f). Scale bars are in micrometers. (a) An array representative of the five out of six control arrays that exhibited no post-aging damage to PPX-C. (b) One control array aged at 87 °C did show almost imperceptible cracking, not visible with backscatter imaging (left of dashed line) but apparent with secondary electron imaging (right of dashed line) after charging from EDS measurements (not colored to show contrast change). (c) Arrays aged in RAA at 67 °C consistently showed cracking of PPX-C. (d) Tip detail showing cracking and evidence of PPX-C removal. (e) One array showed cracking and delamination of IrO_x_ metal. (f) Detail image of PPX-C cracks. Colored images: red – parylene C; blue-green – silicon; light gray – IrO_x_; yellow – silicone. For uncolored images see Fig. S3.

In contrast to controls, the addition of H_2_O_2_ to PBS aging at 67 °C was associated with PPX-C crack formation in all six out of six UEAs (Fig. 4d-f). The damage manifested with differences in the resulting morphology for individual arrays. This damage varied from sparse cracking that did not seem to penetrate the film (one array), to widespread cracking (three arrays) and even signs of erosion (two arrays). Recall that these devices were aged together, experiencing uniform temperatures and concentrations of ROS. PPX-C cracking for one UEA was accompanied by IrO_x_ delamination (Fig. 4d, PP-X on IrO_x_ peeling away from Si). Close inspection of the film surface for these arrays revealed frequent and variously-sized fissures, but little evidence of general surface roughening as was observed from the cat sciatic explants.

Arrays aged at RAA at 87 °C displayed more variation in endpoint appearances than other cohorts (see Fig. 5; uncolored images: see Fig. S4). Out of 14 total arrays, two displayed moderate cracking, which did not reach the extent seen for most devices RAA-processed at 67 °C. Parylene surface degradation was present for at least three arrays, and appeared as roughening accompanied by obvious thinning in several cases. Fig. 5d shows PPX-C cracks and craters not unlike that seen for the sciatic explant (Fig. 2d). Five arrays showed large-area PPX-C removal, also similar to the sciatic explant. Curiously, the remaining four arrays in the 87 °C RAA cohort did not show obvious signs of PPX-C damage. Fig. 5e shows surface detail and PPX-C cross section of one such array, revealing moderate topography and the full PPX-C thickness of ∼6 µm. This is in contrast to a device processed concurrently which showed parylene thinning and removal (Fig. 5f). Surface topography for this array was considerably rougher compared to the undamaged device, and cross sectioning revealed more than 75% loss in film thickness compared to the as deposited PPX-C film thickness. Voids in PPX-C were also visible at the parylene-silicon interface.

**Fig. 5.**
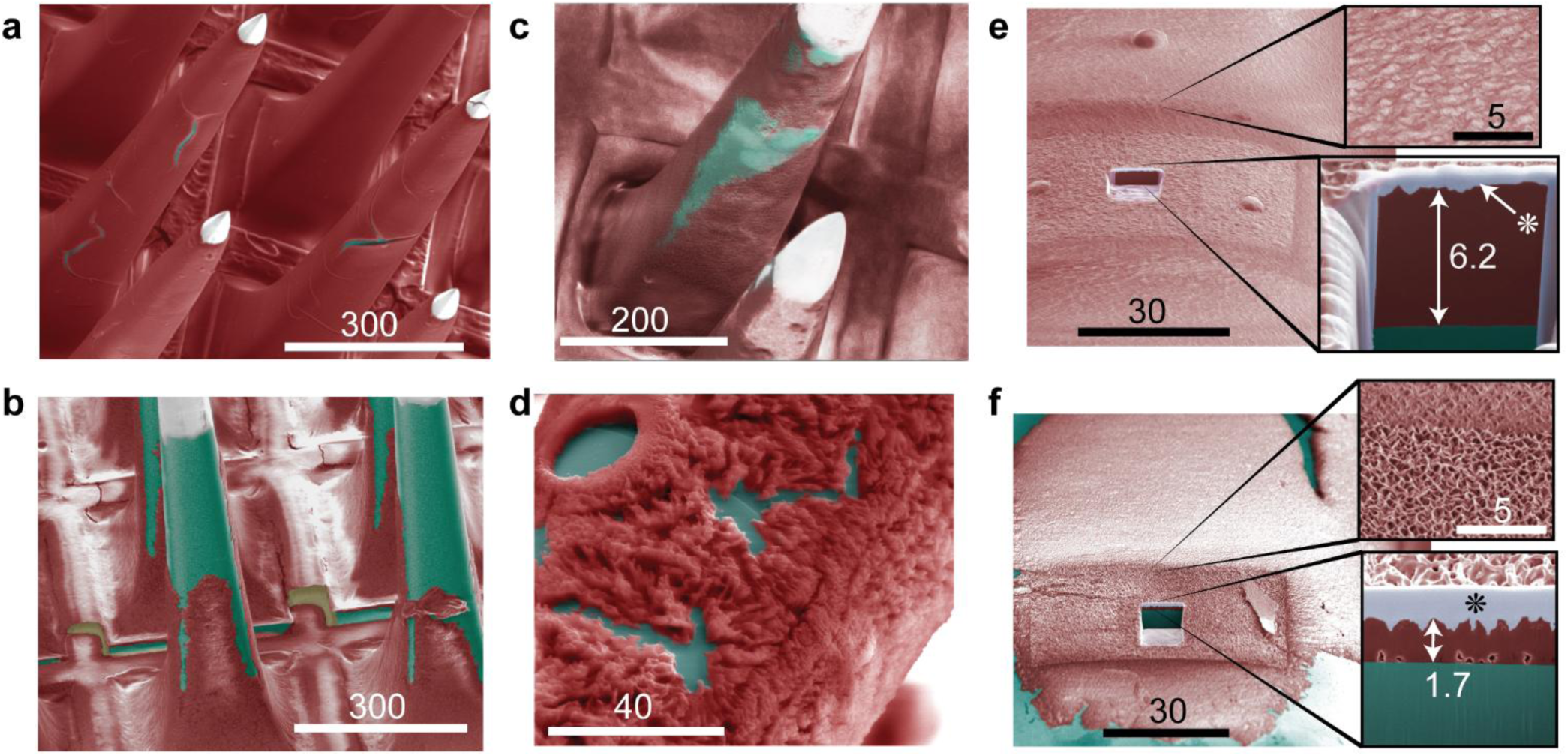
Arrays aged with RAA at 87 °C. Scale bars are in micrometers. Each image is representative of a different array to illustrate the diverse results from RAA at 87 °C. Results varied from (a) moderate PPX-C cracking, to complete degradation and removal: (b) shows erosion presumably originating at the tip and moving towards the base, and (c) shows bulk thinning and removal. (d) Detail PPX-C damage over silicone, showing cratering and surface degradation similar to observed from *in vivo* arrays. (e) An array that did not show obvious signs of damage displayed moderate surface topography and PPX-C thickness similar to the as-deposited film. (f) An array that showed obvious PPX-C erosion displayed obvious irregular PPX- C topography and thickness reduction by at least 75%. For (e) and (f), light blue (marked with *) on top of parylene (red) is protective Pt deposited *in situ*. Blue-green – silicon; light gray – IrO_x_.

Although considerable and varied degrees of PPX-C damage occurred on UEAs aged under both RAA conditions, test structures aged with arrays did not exhibit the same degree of damage. IDEs aged in RAA at 87 °C displayed moderate blistering and delamination on approximately 20% of PPX-C area, but no signs of cracking or surface damage. Similarly, no cracking or surface damage was seen on shafts of t- UEAs (UEAs with complete PPX-C coverage, *i.e.* no tip deinsulation) aged in any condition.

Prior reports from our lab have never noted PPX-C degradation on IDEs aged in PBS at a variety of temperatures [23], [13], nor on fully encapsulated (*i.e.* tips not exposed) UEAs [14]. The primary difference for t-UEA processing is the lack of exposure to O_2_ plasma as part of the tip deinsulation process. These results suggest that oxygen plasma deinsulation of UEAs may play a role in reducing PPX-C robustness, but other factors such as the temperature during the process, and other mechanisms, have not been systematically studied or ruled out.

### 3.2 Electrochemical impedance spectroscopy (EIS)

EIS provided quantitative evaluation of electrode damage, complementing the qualitative characterization of electron microscopy. Impedance measurements for explanted arrays were limited to data taken at 1000 Hz, restricting comparisons between explants and *in vitro* devices. Sciatic and femoral arrays showed pre-implant average moduli of 190 and 263 kΩ, respectively. After implantation, impedance for these arrays increased respectively to 280 and 565 kΩ, and averaged 700 and 1700 kΩ prior to explantation at the 3.5-year time point. Physical characterization of the explants suggests the impedance changes for these arrays were primarily driven by damage to electrode metal. This is further supported by the fact that the impedances increased with time during implantation, which is more consistent with metal degradation than encapsulation failure and its associated impedance decreases. It is clear that degradation of both metallization and encapsulation has occurred. This highlights the complexity in tracking degradation of the electrodes through interpretation of impedance data at a single frequency, though we have found wide band impedance measurements do improve characterization of degradation [14], [17], [47]. The electrode metal deposition process has undergone changes since the fabrication of these implanted arrays, and these modifications have decreased tip impedance mean and variance, as well as improved metal robustness. This improved metal was employed on our test arrays, which exhibited impedances almost two orders of magnitude lower than implants (3-8 kΩ at 1000 Hz), as well as very little metal damage during aging.

In contrast to explant impedance, the impedance spectra for devices aged *in vitro* generally correlated with the extent of PPX-C damage. Representative examples of undamaged and damaged UEA impedance spectra are given in supplemental data (Fig. S5). Control arrays aged at 67 °C, for which no damage was observed, exhibited no more than 10% change to impedance. This is consistent with previous reports of UEA impedance undergoing little change after aging in PBS at 67 °C for durations in excess of 100 days [14]. Most UEAs subjected to RAA exhibited impedance reduction, as has been previously reported for devices aged using RAA [17]. We observed 30% impedance reduction at 100 kHz for devices with slight PPX-C damage, *e.g.* cracking. This reduction did not become more pronounced for instances of increased PPX-C damage. Rather, increased damage was associated with reduced impedance at low frequencies, with heavily damaged PPX-C associated with a 70% reduction at 1 Hz.

### 3.3 Energy dispersive X-ray spectroscopy (EDS)

EDS was used to measure the atomic composition of the films. These measurements indicated that oxygen concentration increased due to aging *in vivo* and *in vitro*, as well as from elevated (>67 °C) temperature processes such as thermal oxidation, plasma etching and deinsulation. The concentration of oxygen in PPX-C was of particular interest, as physical damage to PPX-C and reduction of the impedance was correlated with RAA. Fig. 6a-d shows means and standard deviations of oxygen normalized to carbon (O_norm_) for four different device groups, carefully collected from areas free of organic tissue. Our decision to normalize to the carbon concentration was based on the premise that most oxygen incorporated into PPX-C film would be due to carbon oxidation, as opposed to diffusion of atomic oxygen or film hydration (supported by XPS and FTIR results, discussed in the sections that follow). Results for *in vivo*, control, and RAA arrays are shown in Fig. 6a. Residual PPX-C on *in vivo* arrays exhibited O_norm_ levels of 0.08±0.03, approximately 2× that of 87 °C RAA devices, the next closest group. RAA at both 67 °C and 87 °C was accommodated by >20% increases to O_norm_ compared to respective controls, however this change falls entirely within the standard deviations of both device sets and may be attributed to measurement noise. A more striking trend was observed between control and experimental arrays aged at 67 °C and 87 °C, where O_norm_ levels associated with 87 °C processing increased by >70% over devices processed at 67 °C.

**Fig. 6.**
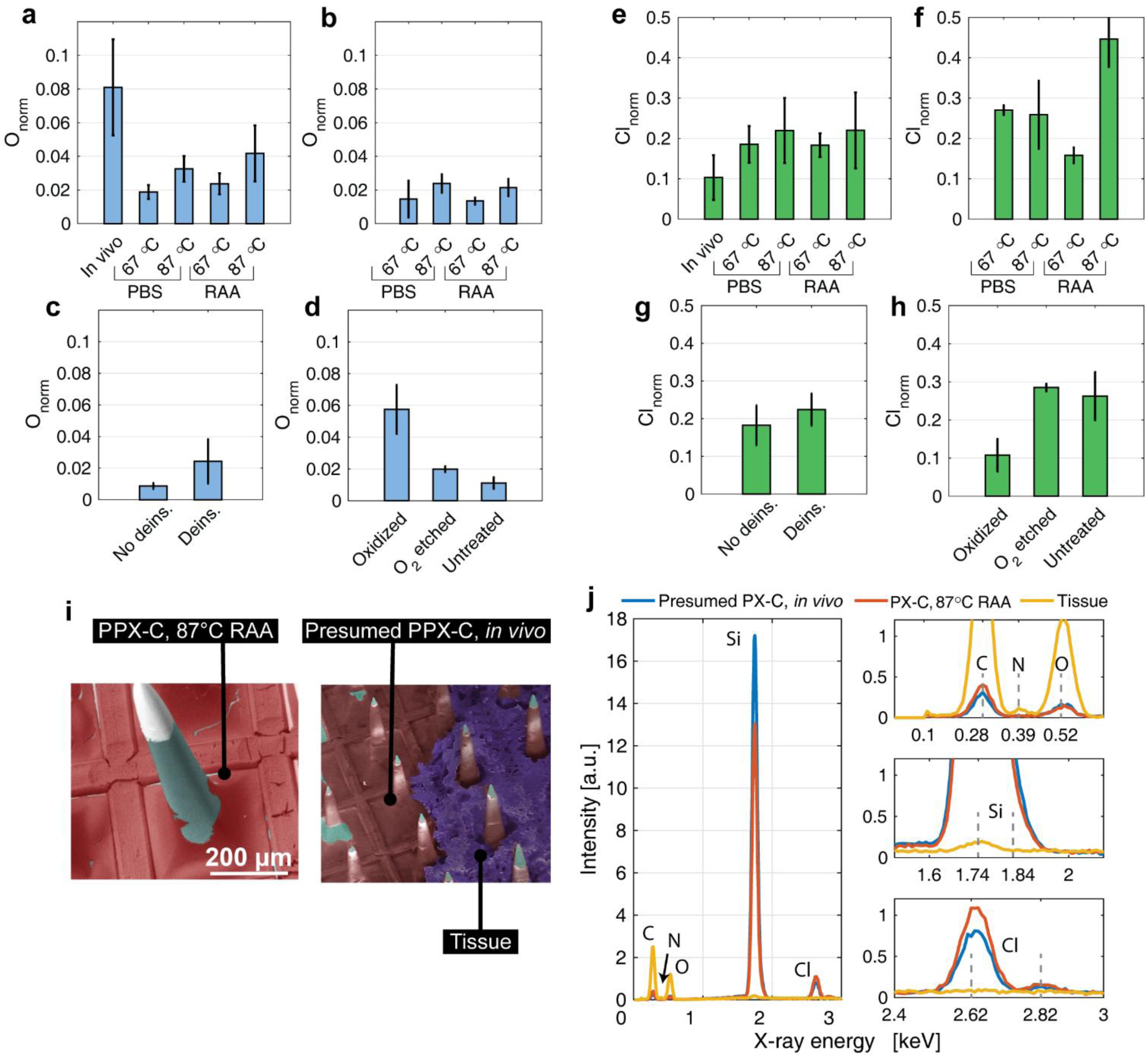
EDS was used for semi-quantitative characterization of PPX-C elemental composition. (a)-(d) Oxygen concentration normalized to carbon concentration (O_norm_) for (a) explant, control, and RAA arrays, (b) test structures (IDEs and t-UEAs), (c) reference arrays before and after oxygen plasma processing, and (d) planar reference samples. Bar graphs show mean and standard deviation. (e)-(h) Chlorine concentration normalized to carbon concentration (Cl_norm_), shown in the same order as O_norm_. (i) Markers illustrate where EDS spectra were taken for data in (j), which shows EDS intensity of PPX-C on an 87 °C RAA array, presumed PPX-C on the sciatic explant, and organic tissue on the sciatic explant. High-resolution plots show details for detected peaks, vertical dotted lines indicate associated X-ray energies used for peak identification.

Measurements of test structures followed similar trends as arrays (Fig. 6b), with samples that were RAA-processed at 87 °C showing on average 60% increase in O_norm_ compared to samples at 67 °C. Test structure controls aged in PBS at 87 °C did not meaningfully differ in O_norm_ from those of the RAA experimental group at the same temperature. Interestingly, O_norm_ values of 0.01-0.02 for test structures were 25-50% lower than those of arrays aged using respectively identical *in vitro* conditions.

One key difference between aged UEAs and aged test structures was that the deinsulation process was performed for the former, but not the latter. As PPX-C damage and elevated O_norm_ were associated with both deinsulated and aged devices, we conducted measurements of reference arrays without and with deinsulation to obtain preliminary data on the effect of deinsulation on the composition of PPX-C. Fig. 6c displays these measurements, which suggest an increase in O_norm_ following deinsulation. One UEA measured with EDS before and after deinsulation showed a very distinct increase in O_norm_, but for other devices no significant change was observed. This is reflected by the coefficient of variation (CoV) of 58% for deinsulated devices.

Planar reference samples subjected to known oxidative damage mechanisms were characterized to assist our understanding of how *in vivo* and RAA damage might be occurring. EDS of PPX-C on polished silicon substrates subjected to thermal oxidation, oxygen plasma etching, or no treatment (control) are presented in Fig. 6d. Thermal oxidation has been reported to be associated with carbonyl bond formation on aliphatic parylene linkages, leading to chain scission [34], [55], and average O_norm_ for these samples was approximately 0.06±0.02, which was between measurements for arrays aged *in vivo* and in 87 °C RAA. Oxygen plasma damage has been suggested to occur via benzene ring opening [53], [56], and yielded O_norm_ of 0.02±0.002, which was closest to control arrays aged at 67 °C and 75% larger than O_norm_ of untreated PPX-C. An increase in EDS-measured oxygen content after oxygen plasma exposure is consistent with previous work [57].

Evaluation of the chlorine concentration normalized to that of carbon (Cl_norm_) is shown in Fig. 6e-h. While stoichiometric PPX-C is characterized by a Cl_norm_ of 0.125, Cl_norm_ calculated from measurements of pristine PPX-C taken from non-deinsulated UEA and planar untreated reference samples was 0.18±0.05 and 0.26±0.06, respectively. This discrepancy underscores the semi-quantitative nature of EDS, and the Cl_norm_ range of 0.18-0.26 was taken as representing the unaged and undamaged condition against which all other devices were compared. *In vivo* samples showed Cl_norm_ values below this range (0.10±0.06), while arrays aged in PBS and per the RAA protocol exhibited Cl_norm_ between 0.18 and 0.22, with no temperature or H_2_O_2_-dependent trends observed. These results suggest that the films were adequately rinsed to remove salts from the physiological solution, which could serve to increase the Cl concentration, which was further supported by the lack of crystalline salt deposits from SEM observations. Measurements of test structure controls aged in PBS also fell within the range of unaged devices. However, RAA-processed test structures showed Cl_norm_ values in excess of 0.26, perhaps due to insufficient device cleaning after aging. Cl_norm_ for thermally oxidized planar references was similar to that of UEAs aged *in vivo*.

While conducting EDS and other spectroscopic measurements, there was concern that the film on *in vivo* arrays that was assumed to be PPX-C was in fact residual organic material. To increase confidence that the *in vivo* measurement we were taking were in fact of PPX-C, we compared EDS spectra of the parylene film in question to known parylene and tissue residue. Fig. 6e indicates where measurements were made for comparison. An array aged with RAA at 87 °C was chosen as a parylene reference due to the presence of RAA-induced thinning and surface damage similar to that seen on the sciatic explant. The organic reference signal was taken from the residual organic material on the sciatic explant, and the *in vivo* parylene signal was taken from presumed PPX-C film on the same explant. Fig. 6f shows all raw spectra overlaid. Considerable similarity was observed between the known and presumed PPX-C films, made more apparent in the detailed spectra to the right of the full spectrum plot. The chlorine peak in particular provided a telling indication of PPX-C film, as it was completely absent from the spectrum of the known biological material. From these data, we are confident that measurements taken from *in vivo* arrays are in fact of PPX-C and not tissue residue.

### 3.4 X-ray photoelectron spectroscopy (XPS)

Compared to EDS, XPS has much greater surface sensitivity as well as the ability to evaluate chemical bonds. While several EDS measurements could be made at different sites for the same array/device, XPS measurements required sampling a larger surface area, such that only one independent measurement was possible for each sample. As received measurements were first collected, followed by sputter cleaning with an argon ion beam to remove surface contamination and extended for depth profiling. This permitted multiple measurements across the same area, but also likely altered the PPX-C composition and chemistry of the film reported by the measurements. An indication of measurement repeatability was obtained by duplicate analysis of four samples (one *in vivo*, two 87 °C RAA arrays, one 87 °C PBS control array), without argon sputtering. CoVs for O_norm_ and Cl_norm_ did not exceed 40% for all devices, and were commonly 15-25%, with inter-device trends largely conserved across measurements. Analysis of carbon binding to oxygen was less repeatable, with one sample showing a CoV of 80%. Therefore, for improved confidence as well as ease of comparison to EDS data, most XPS data analysis was focused on elemental composition measurements. These data are shown in Fig. 7, for which measurements are grouped by aged arrays (Fig. 7a, d), reference arrays (Fig. 7b, e), and planar references (Fig. 7c, f). Regarding sample counts (see Table S1), the repeated measurements previously mentioned were averaged to give an N of 1 for each sample analyzed in duplicate. These means were then averaged with measurements of other samples from matching cohorts to give cohort means and standard deviations, for sample groups of N>1.

**Fig. 7.**
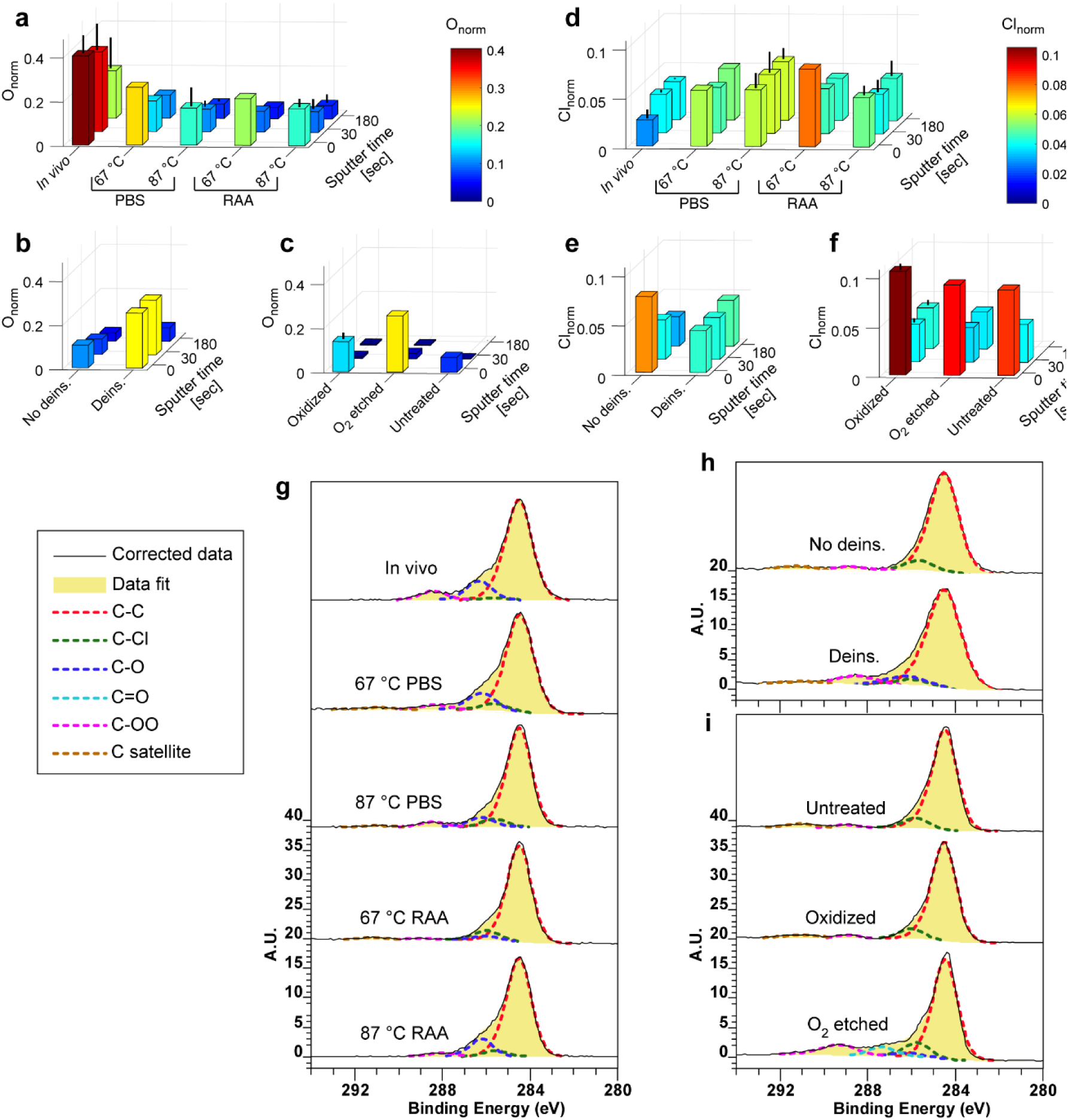
XPS data are shown as concentration (at%) of O (a)-(c) and Cl (d)-(f) for different devices normalized to C. Bar colors correspond to normalized values as shown by color scales. Where present, error bars indicate standard deviations between measurements of different samples. Sputter time refers to argon ion beam sputtering, which removed adventitious carbon, other physiosorbed species (*e.g.* O), and a small amount of the surface material, and enabled depth profiling with longer etches. Region scans of the as received carbon envelope shown in (g)-(i), permitted visualization of high-energy tails associated with carbon bonding to chlorine and oxygen. The legend identifies Gaussian-Lorentzian fits generally in order of lowest to highest binding energy. (a), (d), (g) Aged UEAs; (b), (e), (h) reference UEAs; (c), (f), (i) planar reference samples.

Similar to EDS measures, average as-received XPS measures of O_norm_ for UEAs aged *in vivo* (0.40±0.08) was the highest among all groups, being more than 2× that of arrays RAA-aged at 87 °C (0.17±0.03). We observed little difference in O_norm_ between RAA and control measurements for a given temperature. PPX-C on all aged devices exhibited higher average O_norm_ than pristine PPX-C on non- deinsulated UEA and untreated planar references, which showed O_norm_ = 0.10 and 0.07, respectively, slightly higher than values reported elsewhere [58], [59]. Deinsulated UEA and O_2_ etched planar reference results both showed increased O_norm_ compared to untreated samples, being 0.25 for both, which is consistent with prior characterization of oxygen-plasma etched PPX-C [59]. Also similar to EDS, *in vivo* samples showed the lowest Cl_norm_ of 0.026±0.007.

Unlike EDS results, however, higher aging temperatures were not correlated with higher O_norm_. As-received XPS results indicated the opposite, with aging at 67 °C having higher O_norm_ than aging at 87 °C. In another reversal from EDS trends, higher O_norm_ was measured for oxygen plasma-processed planar reference sample compared to thermally oxidized references (O_norm_ = 0.14±0.03). Thermally oxidized samples also exhibited the highest Cl_norm_ of 0.10±0.01 according to XPS, while the same samples showed one of the lowest Cl_norm_ according to EDS. Cl_norm_ measurements for untreated and O_2_ plasma-treated reference samples were slightly lower than oxidized references and within 5% of each other, in contrast to Cl_norm_ of non-deinsulated and deinsulated arrays that differed by 44%. The information depth of XPS measurements is ∼5 nm, compared to the ∼1 µm information depth for EDS measurements, which is potentially one difference in the measurements. Argon beam sputtering of samples tended to change the measured compositions, which after sputtering became more similar to untreated PPX-C references. O_norm_ for all devices decreased with sputtering, with array measurements reaching similar values to those of the non-deinsulated UEA, and processed planar references decreasing to similar levels as the untreated planar reference. The *in vivo* array was a notable exception, maintaining higher O_norm_ levels than all other devices at all stages of sputtering. The after-plasma treatment reference array also demonstrated little change and the second largest O_norm_ after 30 seconds of sputtering; however, after 180 seconds it was similar to non-explanted devices. These results suggest that while oxidative changes to PPX-C during chronic implantation may penetrate more deeply, oxidation of PPX-C remaining after *in vitro* aging may be limited to the surface. Plasma processing of arrays may incur deeper changes to the PPX-C coating, but this must be verified in future work.

Measurements of Cl_norm_ after 30 seconds of sputtering were near 0.04 for all samples save UEA controls aged at 87 °C, which were closer to 0.06; these values changed little after 180 seconds of sputtering, and are less than 50% of the expected stoichiometry (0.125). This discrepancy may reflect a combination of effects arising from impure PPX-C dimer (known to contain a certain amount of parylene N due to manufacturing limitations) and local chemical changes due to ion bombardment. We note that evidence of Cl^-^ ions was apparent for *in vivo* arrays and UEAs aged at 87 °C (both PBS and RAA), and was accompanied by sodium peaks in the survey scans for these devices both before and after sputtering. Sodium and Cl^-^ were not detected for any other samples, and are potentially attributed to ion permeation of the PPX-C film during aging. Therefore, actual carbon-bonded chlorine content for these devices is expected to be slightly lower than the Cl_norm_ levels reported in Fig. 7d.

Region scans of the carbon envelope between binding energies of 280 and 294 eV had high-energy tails associated with carbon oxidation, as seen for the examples shown in Fig. 7g-i. Spectra are presented with intensity normalized to the C─C peak at ∼284.5 eV. All samples demonstrated indications of ─COO bonds, perhaps due to oxidation of unterminated PPX-C radicals [45], [60] or measurement noise. Intensity of presumed ─COO and ─CO binding was proportional to carbon envelop tail formation, with aged and plasma-processed devices having larger tails and higher oxidation intensities than untreated and thermally processed devices. Larger ─CO intensities were observed for UEAs aged *in vitro* and *in vivo* than for arrays processed in oxygen plasma, while the latter generally had higher ─COO intensity. Evidence of C═O chemistry and a strong carbon envelope tail were seen for the planar O_2_ etched reference, consistent with prior work [59]. Carbon satellite peaks (C-satellite) attributed to benzene ring formation were near the noise limit for most devices, thus their presence was inconclusive.

### 3.5 Fourier-transform infrared spectroscopy (FTIR)

FTIR is sensitive to dipole moments associated with the vibrational characteristics of some chemical bonds, and provided an additional indication of PPX-C chemistry and oxidation, which was clearly evident in spectra for *in vivo* UEAs and thermally oxidized samples. Representative examples of spectra from 3500-670 cm^-1^ (Fig. 8a) had peaks near 1700 cm^-1^ for these devices, attributed to carbonyl bond formation [33], [55], [61]. The remaining samples lacked obvious signs of such oxidation, but shared many other spectral features, a selection of which are identified in Table 3.

**Fig. 8.**
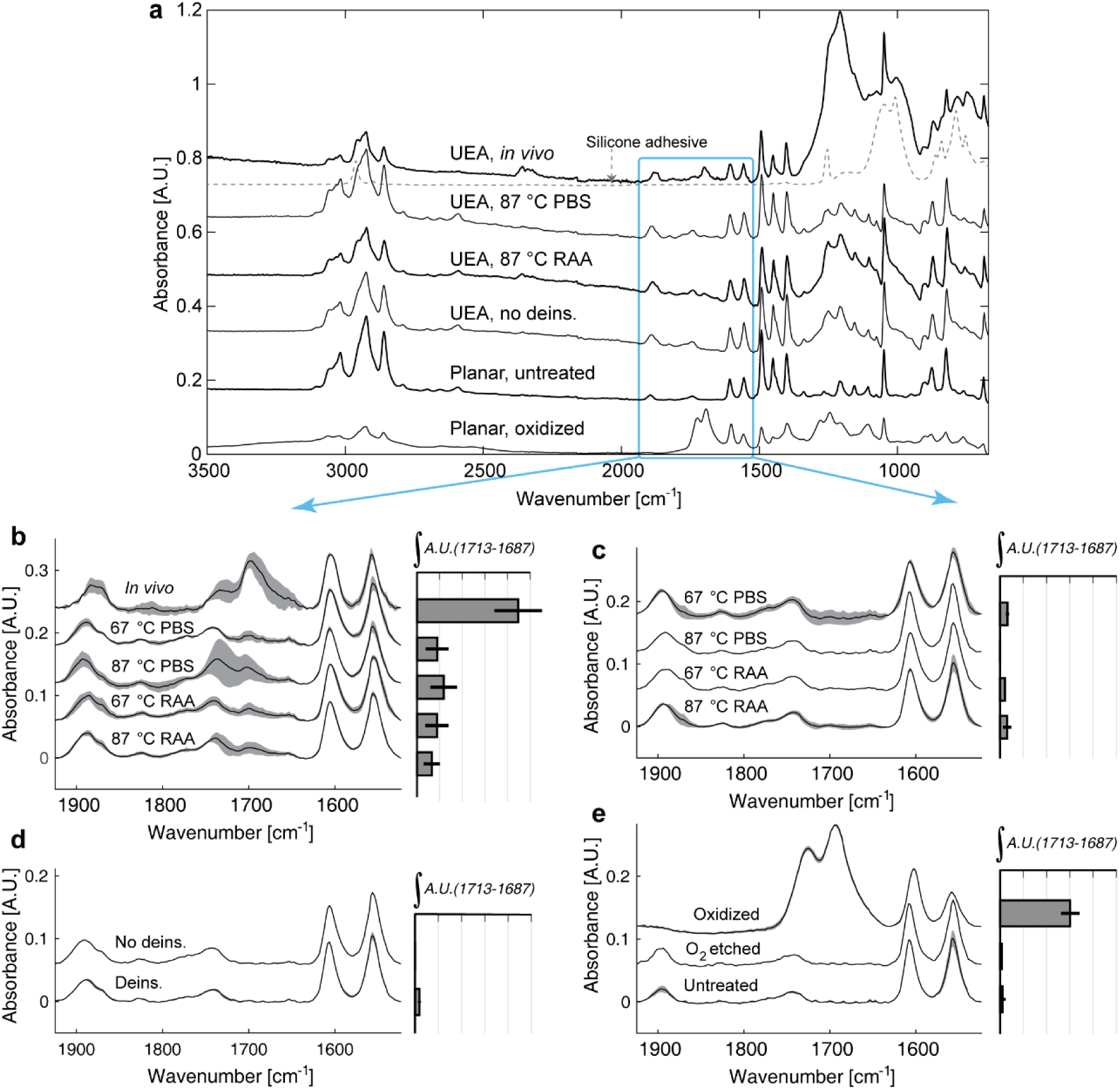
FTIR measurements provided an additional indication of PPX-C oxidation. (a) PPX-C scans from 650-3500 cm^-1^ are shown for representative samples. PPX-C from UEAs was placed on a silicone adhesive, the spectra for which is shown as a dotted line. Background silicone signal was evident for *in vivo* samples due to thinned (<3 µm) parylene film. The region from 1530-1920 cm^-1^ was analyzed for signs of film oxidation for all samples. Averages and standard deviations (where available) of normalized spectra are given for (b) aged UEAs, (c) aged test structures, (d) reference arrays, and (e) planar reference samples. Horizontal bars show area integrated within the window of 1713-1687 cm^-1^, after baselining to window edges.

**Table 3.**
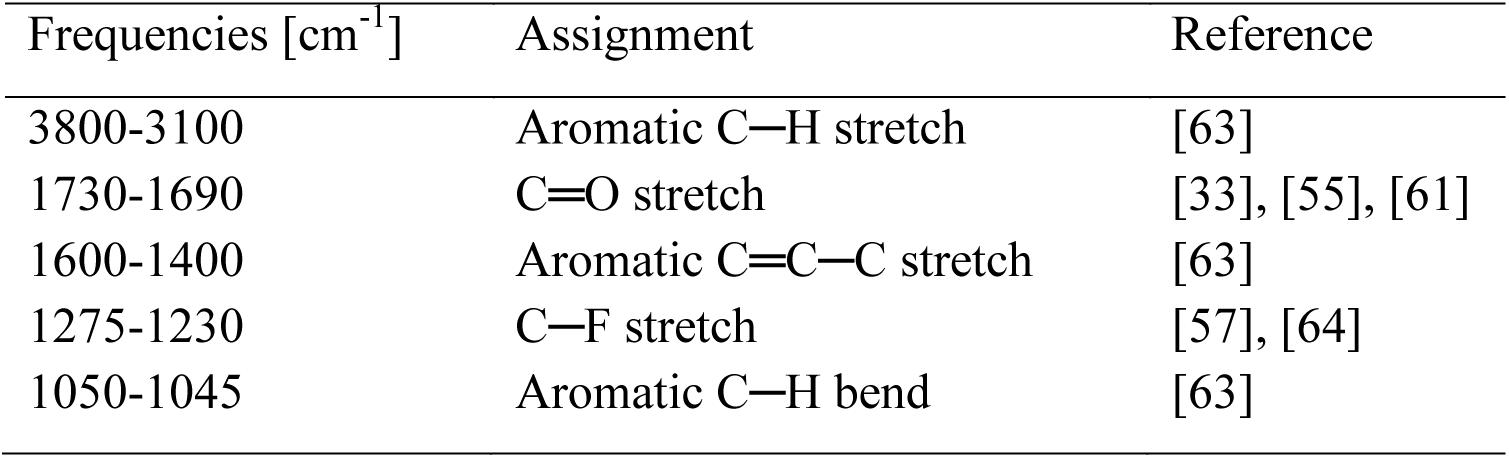
Select FTIR peak assignments for PPX-C spectra.

Pristine PPX-C was represented by the planar untreated condition and was in agreement with previously published infrared absorption data [58]. Spectra of PPX-C from UEAs consistently differed from the planar untreated case due to the inclusion of a broad carbon-fluorine peak near 1250 cm^-1^, a byproduct of the XeF_2_ etching step require to extract PPX-C from the UEAs. This peak was particularly strong for the *in vivo* sample shown and may reflect a higher propensity for C─F reactions in highly damaged PPX-C films. Carbon dioxide absorption peaks near 2350 cm^-1^ for the *in vivo* explants spectrum likely arise from trace carbon dioxide within the FTIR instrument, which was artificially amplified during signal normalization owing to the small signal for *in vivo* explants compared to other samples. Prominent C─F and carbon dioxide features were also observed in the spectrum shown for a UEA heavily damaged by RAA at 87 °C. Virtually no sign of water absorption was noted for aged UEAs, normally indicated by a broad peak formation between 3500-3200 cm^-1^ due to O─H stretching. Since water absorption by as-deposited PPX-C is known to occur and be detectable by FTIR [62], we suspect that exposure of our samples to high vacuum during prior characterization steps resulted in evaporation of a significant amount of the absorbed water.

The silicone adhesive used in mounting UEA PPX-C contributed to FTIR background for some devices, particularly *in vivo* samples due to parylene thickness being <3 µm (similar to or less than FTIR probe depth) for these samples. Therefore, interpretation of spectra at frequencies <1120 cm^-1^ was complicated by the presence of artifact peaks from the silicone. Silicone was the selected adhesive due to a flat FTIR spectrum at frequencies above 1500 cm^-1^, permitting artifact-free comparisons of PPX-C oxidation as interpreted by quantification of the carbonyl peak [33], [55].

To make these comparisons, a two-part piecewise baseline that linearly interpolated between absorbance values at 1925, 1630, and 1524 cm^-1^ was subtracted from each spectrum. Baselined spectra were normalized to the integral of the peak between 1630-1580 cm^-1^, which was largely conserved between all samples, and averaged between cohorts. Averages and standard deviations (where available) of thus-corrected 1925-1524 cm^-1^ spectra for devices are shown in Fig. 8b-e.

As previously noted, oxidation of *in vivo* samples (shown in Fig. 8b) was correlated with a peak at 1697 cm^-1^, between prior reported carbonyl peak values for PPX-C of 1701 cm^-1^ [61] and 1695 cm^-1^ [33], [55]. Detailed observation of UEAs aged *in vitro* (RAA and PBS controls) also revealed signs of carbonyl peak formation, to a similar extent between all *in vitro* samples regardless of processing temperature or condition. The large standard deviation for PBS controls at 87 °C was caused by a sample with a local baseline offset, but which otherwise showed signs of oxidation similar to other samples. Although the presumed carbonyl peaks for *in vitro* UEAs are small, they exist in contrast to most remaining samples. Aged test structures (Fig. 8c) show very slight carbonyl peak formation, if at all, and the same is absent for unaged UEAs (Fig. 8d) as well as all but oxidized planar references (Fig. 8e).

The presence of carbonyl peaks was further quantified by integrating spectral absorbance between 1713 and 1687 cm^-1^, after subtracting an additional baseline linearly interpolated between window boundaries. Horizontal bar graphs in Fig. 8b-e show averages and sample deviations of these integrals for each device group. Peak areas of UEAs aged *in vitro* were over 2× those of test structures so aged, while carbonyl peaks for unaged, nonoxidized samples were smaller still. Quantification of carbonyl peaks for *in vivo* UEAs and oxidized planar references yielded areas at least 3× larger than UEAs aged *in vitro*. Thus, in this first reported use of FTIR to evaluate neural electrode encapsulation, we found strong signs of film oxidation incurred *in vivo*, as well as indications that *in vitro* processing of UEAs may also introduce oxidation to a degree, similar to EDS and XPS data.

### 3.6 Statistical analysis of EDS and FTIR results

Statistical testing of O_norm_ EDS data and FTIR peak areas between 1713 and 1687 cm^-1^ further verified observed oxidation trends in the data. Testing was not performed on XPS data due to insufficient sample sizes. However, devices were grouped into statistical cohorts according to trends in the spectra that were consistent with XPS results. Due to the lack of consistent trends based on processing condition (PBS versus RAA) or temperature (67 versus 87 °C), all *in vitro*-aged UEAs were combined in a single group, as were all *in vitro*-aged test structures. The remaining groups were comprised of *in vivo* devices, non-deinsulated and deinsulated UEA references, and planar untreated parylene references (six groups total). Welch’s ANOVA found statistical significance (p<0.001) between these groups for both EDS and FTIR data, and Games-Howell post hoc testing found *in vivo* devices to be significantly different from all groups, including *in vitro-*aged UEAs. Aged UEAs did show significantly higher FTIR-measured oxidation levels than all groups aside from *in vivo*, suggesting that *in vitro* accelerated aging may be able to approximate oxidation incurred *in vivo*. However, the significant difference between UEAs aged *in vivo* and *in vitro* emphasizes the need for further work to better understand the differences in chemistry between RAA and *in vivo*, and potentially improve the fidelity of the *in-vitro* methodology to better represent *in vivo* processes.

## 4 Discussion

The failure of implanted neural interfaces has been attributed to neural electrode material damage, including degradation of dielectric encapsulation [24], [27], [39], [65]. Eliminating this failure mode requires knowledge regarding the nature of damage incurred *in vivo*, as well as relevant test methods that can evaluate new electrode designs and materials. To meet these needs, we have characterized silicon micromachined electrode arrays that have been aged *in vivo*, compared to existing and novel *in vitro* accelerated aging paradigms, focusing on how such aging affected PPX-C encapsulation. Using characterization methods never before reported for UEAs, we found strong evidence of oxidation to PPX-C film damaged *in vivo*, and that *in vitro* aging paradigms that include oxidative mechanisms may better represent observed degradation mechanisms.

### 4.1 Characterization of in vivo damage to PPX-C

Two USEAs that were implanted in feline peripheral nerve for more than 3 years had severe damage to PPX-C based on SEM and characterization of the devices after explantation. The degradation included cracking, cratering, thinning, and complete film erosion. Spectroscopic characterization techniques were used to better understand any changes to PPX-C chemistry that may have contributed to this damage. Using EDS, XPS, and FTIR, we found consistent indications of elevated oxygen levels in PPX-C from implanted devices compared to samples subjected to accelerated aging and their controls. We also found evidence using EDS and XPS that the chlorine content was reduced for implanted devices. Similar results were observed for thermally oxidized PPX-C reference samples using EDS, FTIR, and to a more limited extent, XPS. These results strongly suggest that oxidative reactions contribute to *in vivo* damage mechanisms, likely arising from ROS generated by the immune respiratory burst and persistent oxidative stress.

To our knowledge, the post-explant conditions observed here represent the most extreme damage to PPX-C incurred *in vivo* reported to date, although considerable damage to other implanted dielectrics such as silicon oxide has been previously noted [12]. Prior reports of PPX-C damage to implanted neural electrodes have found cracking, thinning, and cratering after 3 years [27], as well as cracking and delamination after shorter time points [24]–[26], [39], although to a lesser extent than observed in this study. These devices were implanted for a relatively long time, and also received some of the most detailed characterization efforts for microelectrodes. Another difference between the UEAs of this study and previously reported devices is implant location. The USEAs examined for this report were implanted in peripheral nerve, compared to cortex for many other reports. Oxidative stress is known to occur in both peripheral nerve and cortex in response to implant injury, and although no direct comparison of ROS production is known to exist in the literature, Christensen *et al.* noted similar inflammation timeframes for peripheral and cortical implants when studying the foreign body response in feline peripheral nerve [66]. The authors also noted that peripheral nerve regeneration occurred at the implant site, in contrast to cell death often noted in cortical implant regions, which has been attributed to oxidative stress [29], [30]. These data do not indicate heightened oxidative stress at peripheral implant sites compared to cortical implants, and in fact may suggest the opposite.

Physical movement may also contribute to implant damage discrepancy, as neural interfaces implanted within hind limb nerve are expected to experience more shearing forces during motion of the peripheral nerve, and potentially stronger tethering forces from large displacements, compared to the micromotion of cortical implants. Physical removal of the PPX-C on neural implants cannot be ruled out, particular for films already weakened by oxidation, which is known to reduce parylene tensile strength [61]. However, peripheral implants are commonly fixed in place using surgical techniques, which can be inadvertently assisted by the formation of fibrous capsules around the implant [66]. Furthermore, physical damage might be nonuniform across different aspects of the device. It could preferentially occur on edge electrodes where external forces might act more directly, compared with a lesser effect on center electrodes where the surrounding tissue is stabilized by the electrodes. Alternatively, there is some evidence of significant vascular compromise near the base of the array, which might allow elevated concentrations of ROS to form. However, no such trend was observed for the explanted devices analyzed here. Fully elucidating the nature of the damage we observed on explanted USEAs, as well as how representative it is of neural electrodes, will require deep analysis of more samples. Such multiyear implantations are difficult experiments, thus characterization of the relatively few devices explanted after such long time frames is of great value. Furthermore, the difficulty of chronic *in vivo* experiments drives the need for accelerated aging testbeds that can more efficiently evaluate new material robustness. Based on our results, the RAA method reported here is promising in that regard.

### 4.2 Characterization of in vitro damage to PPX-C

RAA processing at 67 and 87 °C for durations of 28 and 7 days, respectively, resulted in consistent damage to PPX-C not replicated by aging in PBS alone. These timepoints would represent the equivalent to 224 days at 37 °C based on the commonly used acceleration equation [67][8], [13], but the chemistry of these putatively oxidative reactions might have significantly different kinetics. RAA processing at 67 °C resulted in PPX-C cracking visible in at least 5 out of 6 UEAs. RAA processing at 87 °C induced aging effects including cracking, thinning, cratering, and complete removal, with similar topography as observed from *in vivo* USEAs. In contrast, control samples aged in PBS at the same temperatures yielded very little change to PPX-C observed in SEM, with only one UEA out of more than six processed showing signs of crack formation after aging at 87 °C. This clearly suggests that oxidative degradation is one of the most significant degradation mechanisms for the devices.

We expected that differences in spectroscopic characterization would accompany the observed physical changes between RAA-processed UEAs and PBS controls. Impedance measurements indeed reflected physical damage to PPX-C, and RAA processing was commonly accompanied by impedance decreases throughout the measured spectrum. Impedances for devices aged in PBS changed little compared to RAA cohorts, agreeing with previous work [17]. However, differences attributable to RAA versus PBS processing were less apparent using EDS, XPS, or FTIR. All modalities showed heightened levels of oxygen in aged PPX-C films on UEAs compared to untreated, unaged PPX-C. EDS showed signs of temperature dependence on oxygen concentration, which could be evidence of increased diffusion of water, dissolved gases, or a progression of oxidative reactions into the PPX-C at higher temperatures, but this trend was not repeated with XPS and FTIR measurements. XPS analysis of the carbon envelope and FTIR measurements from 1713-1687 cm^-1^ indicated that oxidation of PPX-C carbon bonds accompanied *in vitro* aging, regardless of H_2_O_2_ presence. This agrees with previous work that found increased oxygen concentration in PPX-C after immersion in 0.9% NaCl and artificial body fluids lacking oxidative constituents beyond water and the associated dissolved oxygen [54].

Strikingly, no physical damage was observed on test structures that included non-deinsulated UEAs and IDE devices, despite simultaneous RAA processing with UEAs that did experience significant degradation. This lack of damage was accompanied by reduced oxidation compared to aged UEAs measured by EDS and FTIR. We cannot attribute this difference to the as-deposited parylene film, since test structures and UEAs were coated with PPX-C within the same deposition run. The most notable differentiating factor between the two device sets was the O_2_ plasma deinsulation process, which is a necessary step for UEA fabrication but was not used on test structures. This suggests that the oxygen plasma deinsulation procedure may play a role in PPX-C aging damage. Similar to UEAs, test structures showed no difference in oxidation level based on presence or absence of H_2_O_2_ during *in vitro* aging. EDS revealed a temperature-dependent trend in O at% as was observed for UEAs, but this could not be confirmed by other modalities.

### 4.3 Effect of the deinsulation process on PPX-C

Observations of as-fabricated arrays before aging did not detect any sign of physical damage to PPX-C after tip deinsulation. We have on occasion observed crack formation in PPX-C on deinsulated UEAs by SEM imaging directly after production, leading to rejection of the devices for quality control. Interestingly, these devices would have likely passed traditional quality metrics such as impedance measurements and optical imaging. The operative assumption has been that PPX-C films that do not undergo visible changes during the deinsulation process have similar integrity to the as-deposited film. However, the spectroscopic characterization reported here of PPX-C on non-deinsulated and deinsulated UEAs suggest otherwise. EDS and XPS characterization found increased oxygen at% on films subject to deinsulation, at levels similar to those seen for deinsulated UEAs aged *in vitro*. FTIR also indicated increased carbonyl signal for PPX-C on deinsulated arrays compared to non-deinsulated UEAs, although this signal was much smaller than those of aged UEAs. In addition, XPS measurements noted loss of chlorine for a deinsulated device compared to a non-deinsulated reference. This information, taken into account with the observations of damaged PPX-C on aged UEAs but not on t-UEAs or IDEs, strongly implies that the UEA deinsulation process can potentially oxidize the PPX-C which reduces resilience against aging. If this degradation mechanism is borne out by additional studies, mitigations strategies to control the temperature excursion and/or dose of reactive oxygen species from the plasma are possible.

PPX-C remaining on UEAs after deinsulation has historically been considered undamaged by oxygen plasma, due to the foil mask designed to protect the body of the encapsulated device. However, differences in PPX-C film chemistry were found using EDS and XPS analysis from before and after deinsulation, and deinsulation processing, which correlated with RAA damage. Preliminary investigations have found temperatures within the plasma chamber can reach in excess of 100 °C. Furthermore, the ambient in the chamber is pure O_2_, and radicals and reactive oxygen species are generated by the plasma. Also, the elevated temperatures may alter the integrity of PPX-C that is masked from plasma by the aluminum foil. Heat treatment of parylene has been studied extensively, and is known to increase film crystallinity [51]. When performed in an inert atmosphere such as nitrogen, annealing and associated crystallinity changes have been noted to increase tensile strength [20], as well as reduce the rate of water diffusion through films [62]. Less advantageously, compromised substrate adhesion has been a noted byproduct arising from thermal mismatch between annealed parylene and its substrate [45], resulting in increased blister formation during soak tests [68].

In contrast, we and others have noted film embrittlement and cracking when parylene is heat-treated in an oxygen-containing atmosphere [45], [69]. The oxygen pressure within the plasma chamber during deinsulation is 0.4 Torr, 0.25% of the oxygen concentration in atmosphere at standard temperature and pressure. Thus a smaller but significant flux of oxygen continues to impinge on the surface. An impingement rate of more than 10^5^ mono-layers per second would be present at this pressure. In addition, reactive oxygen species such as atomic oxygen (neutrals and ions), and ozone have been reported in O_2_ plasmas, which are much relatively strong oxidizers compared to pure O_2_. The high reactivity of oxygen radicals within the active plasma chamber may unpredictably alter reaction kinetics during deinsulation to favor film oxidation. The relative contributions of temperature and gas chemistry will require additional studies.

The oxygen plasma deinsulation process is performed for all UEAs and therefore may be a contributing factor in many reports of PPX-C damage *in vivo* [25], [26], [39], [70]. Schmidt *et al.* noted PPX-C damage on iridium microwires which were deinsulated by exposure to a heated element or high-voltage arcing [27], [71]. In these cases, exposure of PPX-C to extreme heat in oxygen containing environments may have resulted in degradation to the parylene films and facilitated subsequent *in vivo* damage.

The evaluation of deinsulation effects on PPX-C was not a primary focus of this study, thus our non-deinsulated and deinsulated reference sample sizes were small. Due to the nature of sample preparation, EDS, XPS and FTIR analyses were performed on different individual deinsulation samples, and variability in measured oxidation was found between modalities and samples. This variability may reflect variability in the film preparation and deinsulation processes, and/or may play a role in the different damage modes of RAA-processed UEAs. PPX-C film nonuniformity at the micro and macro scale may also play a role, requiring further study of PPX-C characteristics across substrate area, substrates within a single run, and substrates from different runs.

RAA testing of UEAs and test structures has definitively brought to light the possible negative effects of the deinsulation process, and further study utilizing larger sample sizes to understand these effects and mitigate them is warranted. Improved plasma systems or alternative techniques such as laser ablation [72] may be able to decrease oxidation during the deinsulation process. Novel neural electrode technologies incorporating PPX-C are constantly in development, and while common characterization techniques such as EIS have been used to assess heat treatment effects on impedance [73], we have not found impedance to be predictive of PPX-C stability during aging. In such cases, additional spectroscopic methods and RAA can provide deeper insight into possible changes to PPX-C chemistry due to high-temperature processing, as well as an indication of PPX-C resilience to aggressive environments encountered *in vivo*.

### 4.4 Aging damage mechanisms

Little is known about the physiological mechanisms that directly contribute to PPX-C damage *in vivo*. However, considerable effort has been made to understand how thermal and photolytic oxidation influence parylene characteristics, and can help elucidate damage mechanisms at play. Thermal oxidation associated with the appearance of the carbonyl peak between 1701 and 1695 cm^-1^ has been tied to the formation of ester bonds at aliphatic parylene linkages, leading to chain scission upon further reactions with oxygen [33], and film thinning [34]. *In vivo*, ROS produced by the respiratory burst including H_2_O_2_ as well as the superoxide anion •O^-^ and highly reactive hydroxyl radical •OH could initiate hydrogen abstraction at the aliphatic linkage, eventually leading to chain scission and thinning by hydrolysis. Embrittlement and reduction of PPX-C tensile strength through oxidation, as noted for thermally oxidized samples, could lead to crack formation as PPX-C swells from fluid absorption [50]. Photolytic cleavage of chlorine has been identified as another pathway whereby radical sites on PPX-C chains can be created, leading to intramolecular phenylation and hydrogen abstraction [61]. The reduced chlorine measured from explanted devices suggests that a similar pathway may have occurred *in vivo*.

The RAA system was designed to mimic the effects of ROS *in vivo*, but as oxygen and chlorine content of RAA UEAs were different from *in vivo* USEAs and not distinguishable from UEAs aged in PBS, further studies would be required to more definitively indentify the *in vitro* mechanism of damage. PPX-C cracking suggested loss of tensile strength due to oxidation followed by swelling, but cracking was nearly absent for UEAs aged in PBS, which showed similar measurements of oxygen content. We observed thinned PPX-C on several UEAs aged with RAA at 87 °C, and it is possible that analysis of the PPX-C lost to solution, such as through HPLC or mass spectrometry, may yield further clues concerning RAA damage mechanisms. Our choice of characterization modalities was based on prior work that identified similarities between polymers aged *in vivo* and using oxidative test beds, particularly using FTIR [74], [75]. Although RAA damage mechanisms still remain unclear, inclusion of absorption and emission spectroscopic techniques in our study regimen has yielded several novel insights to inform improvements to neural electrode design.

### 4.5 Choice of neural interface characterization modalities

Much work of great value to the field of neural interfaces has been published concerning the characterization of neural implants in the context of *in vitro* and *in vivo* environments. Such characterization has been commonly conducted using impedance spectroscopy and electron microscopy [17], [26], [39], [40], [76], chosen for their applicability to neural electrode performance and ease of use. These techniques are effective for evaluating physical changes to materials, as well as indirect investigation of material chemistry changes inasmuch as these changes are reflected in altered impedances. For UEAs with undamaged IrO_x_ electrode metal, we found impedance reductions correlated with encapsulation damage. These impedance changes were most apparent at different frequencies depending on the extent of PPX-C damage. Mild, severe, and complete PPX-C damage modes were most reflected in impedance changes near 10^5^, 10^3^, and 1 Hz, respectively, underscoring the value of full-spectrum impedance measurements for electrode characterization. The 10^3^ Hz impedance point has been historically used for neural electrode characterization [65], [77], [78] owing to the characteristic frequency of neural action potentials being on the order of 1 kHz, but this single data point does not fully capture the progression of material damage. Agreeing with prior work [17], we affirm the value to neural electrode characterization of capturing impedance data across multiple frequency points.

However, impedance data alone is not sufficient to characterize the electrode material condition, particularly in the case of multiple degradation mechanisms with competing impedance effects. Such was the case for explanted devices, which exhibited increased impedance over time attributed to tip metal damage. This increase in impedance masked any reduction in impedance resulting from encapsulation damage, which was also considerable. Complementing impedance spectroscopy, careful electron microscopy using both secondary and backscattered contrast mechanisms was valuable in identifying multiple physical markers of material damage, including degradation of both electrode metal and encapsulation. Additionally, we employed FIB cross-sectioning to probe the electrodes and their component materials in cross-section, and for the first time found confirming evidence of PPX-C thinning from *in vivo* aging and RAA processing, as well as signs of silicon erosion *in vivo* underneath the IrO_x_ tip metallization. These findings aid our understanding of the damage modes and the challenges faced by implanted neural electrodes, and underscore the value of cross-section analysis in addition to SEM surface imaging for investigating the underlying degradation modes.

Not surprisingly, pre-aging characterization using SEM and EIS showed no predictive power for the occurrence of device aging damage, and post-aging characterization with these modalities provided very little information concerning the nature of such damage. By utilizing EDS, XPS, and FTIR we found the first clear signs of oxidation *in vivo* as well as *in vitro* for neural electrodes, and identified processing steps which may have an impact on PPX-C robustness. The utilization of these methods in neural electrode characterization will greatly aid informed design of future electrodes to identify and mitigate failure modes. However, obstacles such as sample preparation and the potential for damage to the device might limit their use. Having a method to detect susceptibility to oxidative degradation would be of high utility. Further effort would be needed to determine if EDS or FTIR would be possibilities, and justify the time and expense as quality control techniques.

Of the three utilized characterization techniques, EDS is the simplest to employ and has indeed been used previously to characterize explanted microwire electrodes [31]. The technique is considered semi-quantitative, and therefore, care must be used in evaluating trends in the data. XPS can be used to complement EDS measurements, but requires extensive time, expertise, and expense to perform, as well as creative sample preparation or analysis techniques for devices with considerable topography, such as the UEA. In addition, discrepancies between EDS and XPS such as those we observed between thermally oxidized and oxygen plasma-etched planar references may arise from the surface-sensitive nature of XPS. FTIR is not as surface sensitive and is widely used for characterizing polymer films, but the volume of analyte required for reliable FTIR analysis can restrict its use. Prior work has evaluated PPX-C for neural microelectrode applications utilizing FTIR, but the small size of actual implanted devices limited FTIR characterization to planar monitor samples soaked *in vitro* [15]. However, we found that the volume of parylene removed from a 4x4 UEA was more than sufficient for repeatable ATR-FTIR measurements. Despite the challenges posed, utilization of characterization methods such as these, in addition to electrochemical and electron microscopy techniques, will greatly enhance neural electrode lifetime data and drive improvements to informed electrode design.

### 4.6 Future work

In this work, we have found favorable comparisons between UEAs aged *in vivo* and using RAA. However, the small sample size of the explanted device cohort limits the confidence of our conclusions, and the strength of our comparisons with previous reports of PPX-C aging is weakened by the fact that most electrodes of previous reports were used in recording applications, while the USEAs of this report were largely used for stimulation. Therefore, the impact of electrode function on PPX-C damage must be addressed in future work, and analyses of more explanted devices is needed to improve knowledge of conditions encountered by neural microelectrodes, and engineer solutions.

Additional characterization methods can also provide more insight into the changes incurred *in vitro* and *in vivo*. While crystallinity and tensile strength have been noted to be affected by oxidation, our characterization techniques were not suited to such measurements, and performing the measurements on explanted devices would be challenging. Preliminary work found that differences in PPX-C Young’s modulus based on presence or absence of oxidation could be detected using nanoindentation and peak force tapping atomic force microscopy (Bruker Dimension ICON, Santa Barbara, CA); however, our efforts to apply these modalities to UEAs were unsuccessful. Through the use of additional methods such as these, the degradation mechanisms may be better elucidated.

The RAA system was designed as a test bed that subjects devices to a presumed worst-case scenario *in vivo* incorporating ROS constituents. Such systems have been previously investigated for testing polymers, *e.g.* environmental stress cracking of polyurethanes. Meijs *et al.* found effects similar to *in vivo* stress cracking when polymers were exposed to H_2_O_2_ at 100 °C [79]. Zhao *et al.* found similar SEM and FTIR characteristics between devices aged *in vivo* and *in vitro* when the *in vitro* system included exposure to H_2_O_2_ in addition to blood plasma proteins [74] or cobalt chloride [75]. The latter system was designed to produce the •OH through Fenton chemistry, known to occur *in vivo* as part of the respiratory burst [80]. The production of •OH through interactions between H_2_O_2_ and iron cations has also been associated with parylene cracking in physiological electrolytes at room temperature [81]. These studies present the possibility of modifying the RAA system to include additional reactants known to drive oxidation processes, with which improved neural electrode devices and material choices can be evaluated.

## 5. Conclusion

In order for neural microelectrodes to be a clinically viable technology, material damage from chronic exposure to the physiological environment must be prevented. This requires understanding damage mechanisms and having suitable test protocols for evaluating potential solutions. However, knowledge in these areas is lacking for neural microelectrode interface technologies, for which utilization of traditional and well-vetted materials is currently not tractable. Mechanisms and solutions are specific to the type of material and its function, and in this work we chose to better understand the damage mechanisms of PPX-C, a widely-used and well-regarded dielectric insulating film for neural interfaces. We also observed notable degradation in the IrOx tip metallization and Si of the shank, though these damage modes are not the focus of this work. Prior work has shown evidence of PPX-C damage from *in vivo* exposure using SEM and EIS [24], [27], [40], [76], but these modalities did not provide information concerning chemical changes that may point to damage mechanisms and potential solutions.

In the present work, we identified some of the most extensive damage appreciated to date from a unique population of Utah Slanted Electrode Arrays implanted in the sciatic peripheral nerve for 3.25 years. These observations contrast with the widely-held favorable impressions for PPX-C, and therefore warranted further investigation. We have developed methods to analyze PPX-C film chemistry from explanted USEAs using absorption and emission spectroscopy techniques, and found evidence of film oxidation and chlorine abstraction for these films. We replicated aspects of *in vivo* PPX-C degradation through oxidative RAA, the first reported time such damage has been recreated *in vitro* for neural interfaces. However, while UEAs aged *in vitro* showed signs of oxidation compared to unaged UEAs, overall PPX-C film chemistry for devices aged *in vitro* was different from devices aged *in vivo*. These results suggest that RAA is an important soak testing paradigm, as it produced much more representative degradation of PPX-C compared to our extensive prior saline soak testing efforts, which never generated degradation of the caliber seen in RAA and long-term *in vivo* samples. This is supported in our case because using RAA enabled us to identify oxygen plasma processing of UEAs as a potential cause underlying PPX-C damage *in vitro* and even *in vivo*. This underscores the value of such testing for identifying possible material failure modes. This suggests RAA is complimentary and superior to typically employed *in vitro* aging systems comprised solely of buffered saline solution at physiological temperatures or higher, which historically have not uncovered device failure modes prior to costly *in vivo* application and testing. There is a clear need, however, for further work to better understand material aging mechanisms *in vivo*, how such mechanisms may be simulated in an accelerated fashion *in vitro*, and their kinetics. Such will permit quantitative lifetime predictions for further materials optimization. We noted that because of the more complex chemistry of the oxidative reactions, a wider range of acceleration factors are likely, highlighting the need for greater care in the collection and interpretation of results from such studies.

Future work to meet this need must include more in-depth study of devices explanted after chronic *in vivo* use, including chemical spectroscopy modalities such as those presented here. The sample size of two explanted USEAs employed in this work was sufficient to demonstrate the value of new characterization techniques and compare RAA outcomes, but general conclusions regarding *in vivo* aging cannot be made with confidence using such a small sample set. As these devices were implanted in peripheral nerve and used for stimulation, differences in material aging may exist with devices implanted in cortex and used for recording. Further work must also incorporate careful material characterization prior to implantation/aging to place post-aging characterization in context, as illustrated by our finding with regard to PPX-C on deinsulated UEAs.

Building on work done using the RAA system, many opportunities exist to further develop *in vitro* aging techniques, such as incorporating acidic species to further mimic the foreign body response. As previously mentioned, stimulating neural electrodes may undergo different damage modes from recording electrodes, giving value to test methods which combine stimulation with soak testing. Importantly, better understanding of activation energies for materials and test methods is needed to confer greater accuracy to equivalent durations of accelerated aging. Common practice is to assume a doubling of aging rate for every 10 °C increase in aging temperature [8], but we found that RAA at 67 and 87 °C caused different damage modes, despite adjusting aging time according to this relationship. This suggests that the current assumptions for accelerated aging are not suitable for all situations without proper justification, and future development of such tests should include studies of reaction kinetics to provide such justification. Through such testing and analysis, data-driven design of medical devices and material choices will facilitate improved outcomes and clinical adoption of novel technologies to improve patient quality of life.

## 6 Acknowledgements

The authors would like to thank Dr. David Warren for assistance with *in vivo* sample preparation, Dr. Paulo Perez for assistance with XPS analysis, and Dr. Nicholas Ashton for FTIR guidance. Funding for this work was provided in part by the NSF through IGERT program number 0903715, and by the NIH through grant 1R43EB018200-01A1 and the DARPA HAPTIX program (N66001-15-C-4017), as well as the DARPA Inter Agency Agreement with U.S. Food and Drug Administration.

The views, opinions, and/or findings contained in this article are those of the authors and should not be interpreted as representing the official views or policies of the Department of Defense or the U.S. government. The findings and conclusions in this paper have not been formally disseminated by the Food and Drug Administration and should not be construed to represent any agency determination or policy. The mention of commercial products, their sources, or their use in connection with material reported herein is not to be construed as either actual or implied endorsement of such products by the Department of Health and Human Services.

## Data availability

Raw and processed data and images are available upon request from the corresponding author.

---

**Fig. S1.**
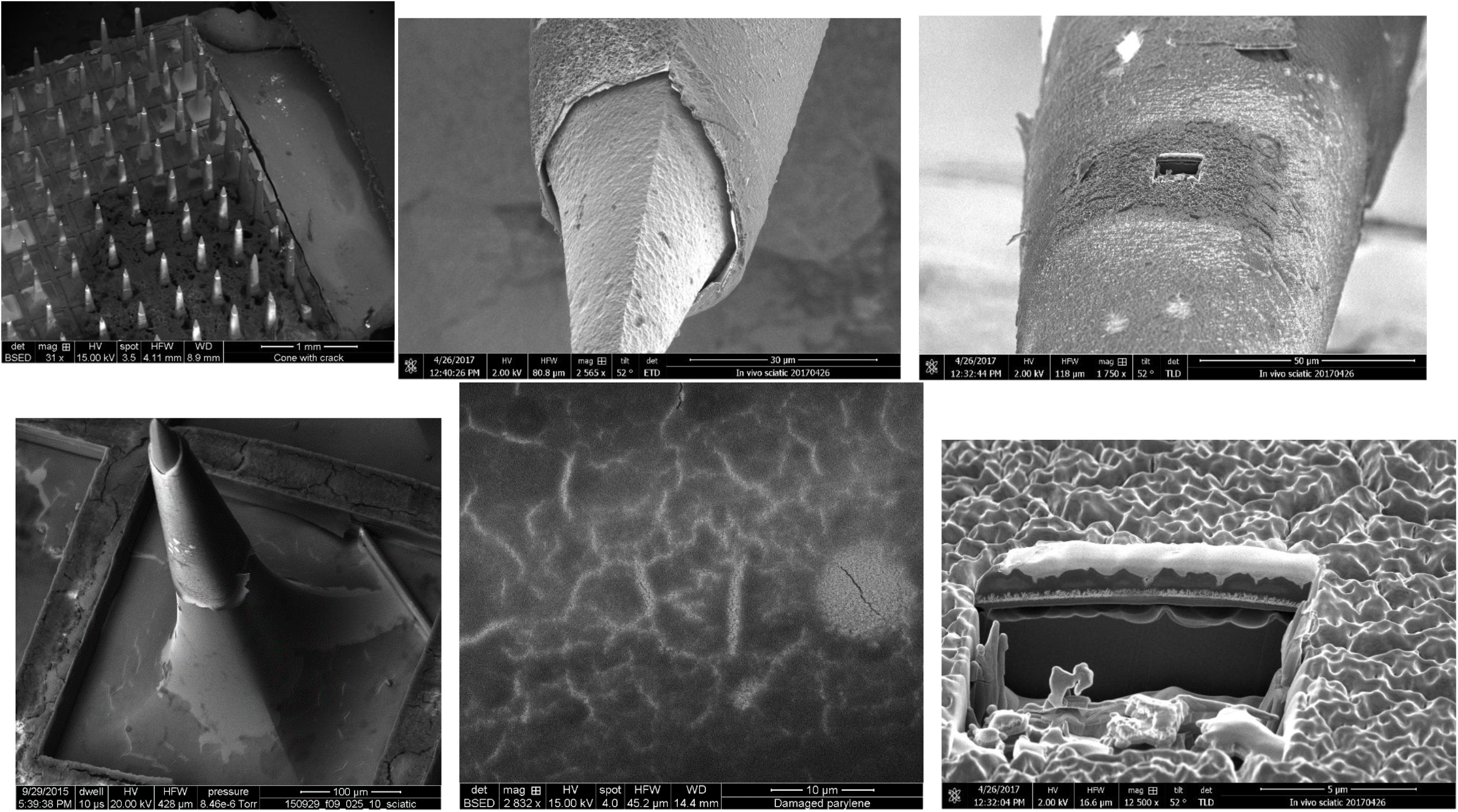
Select raw images used for Fig 2

**Fig. S2.**
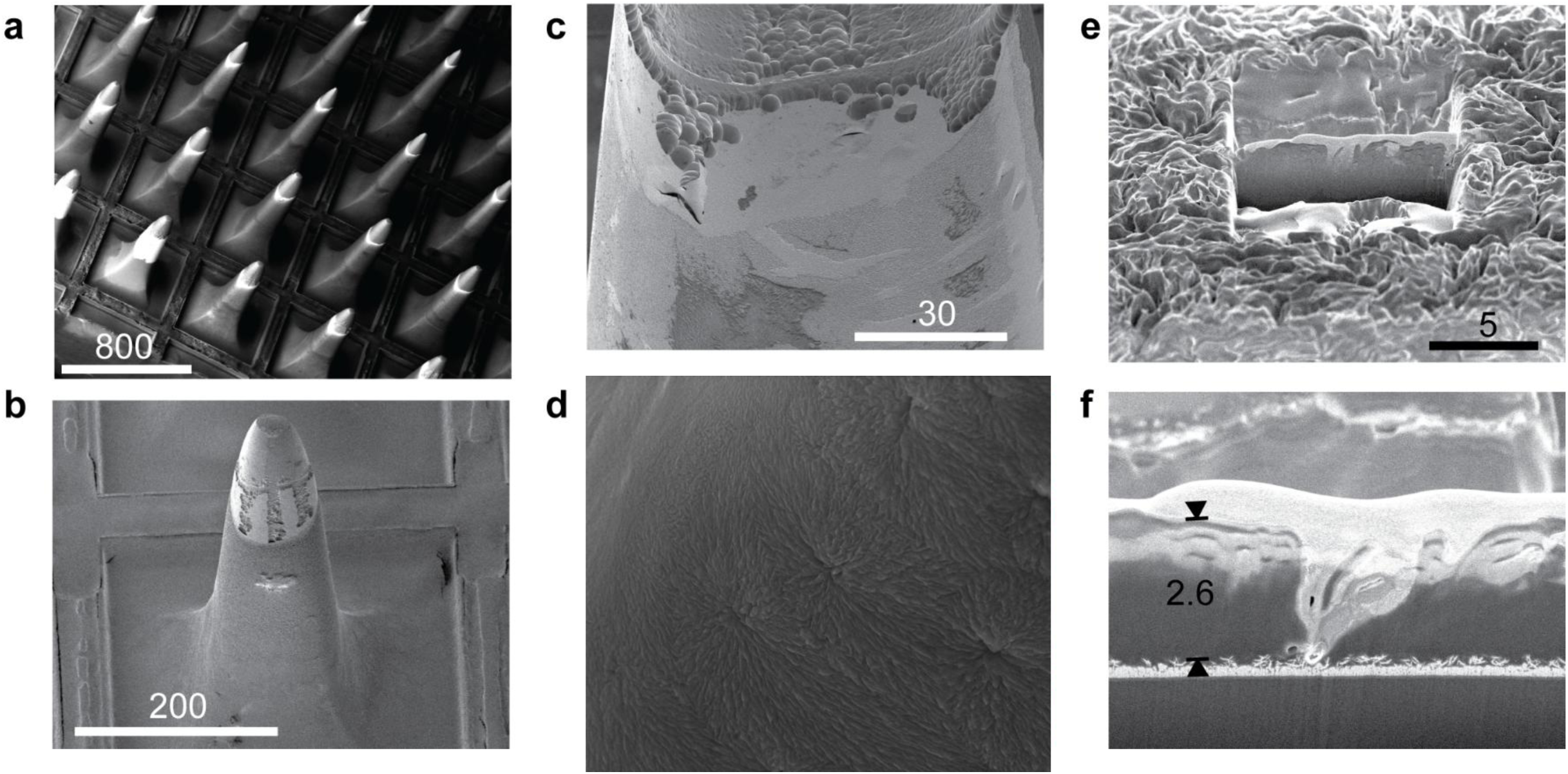
Select uncolored images used for Fig 3

**Fig. S3.**
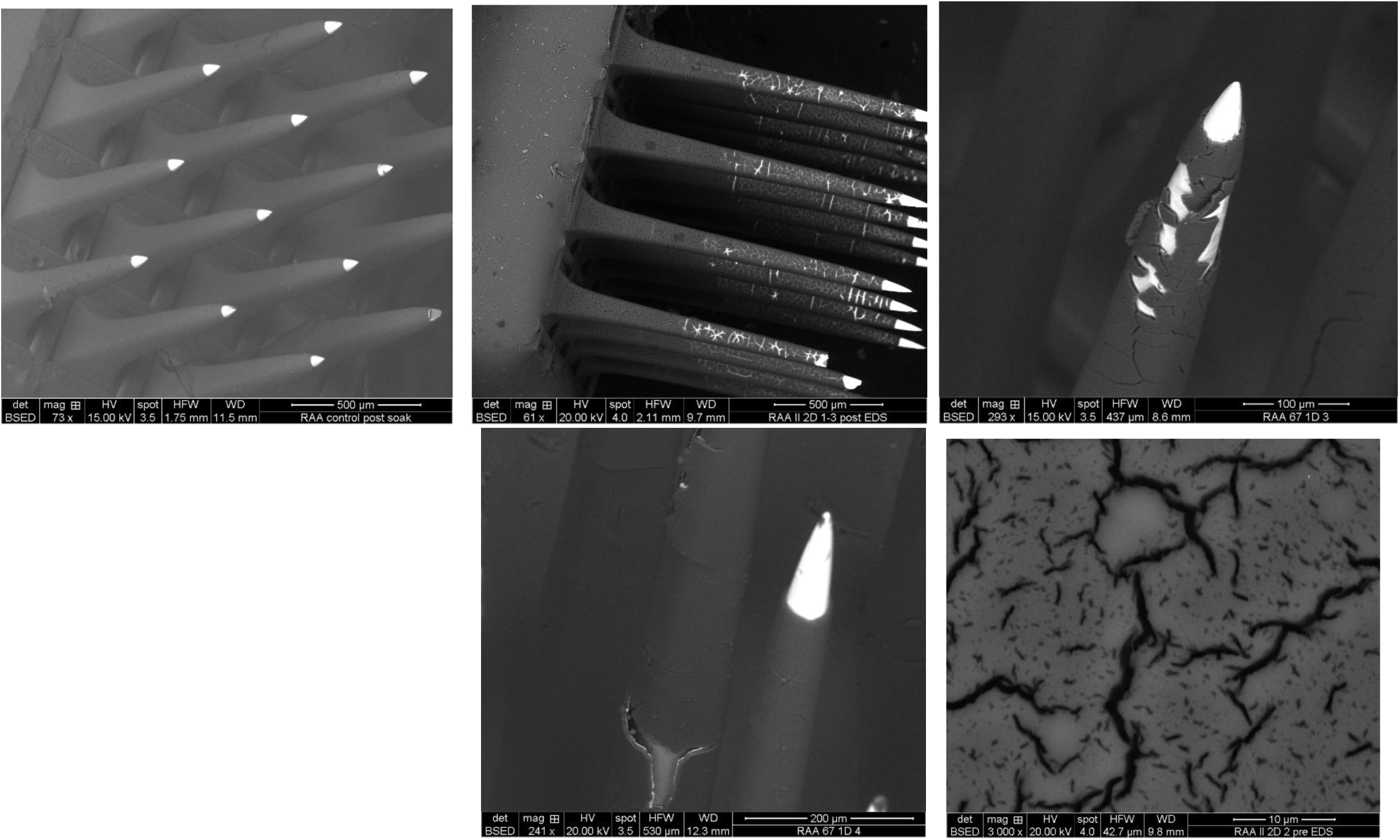
Select raw images used for Fig 4

**Fig. S4.**
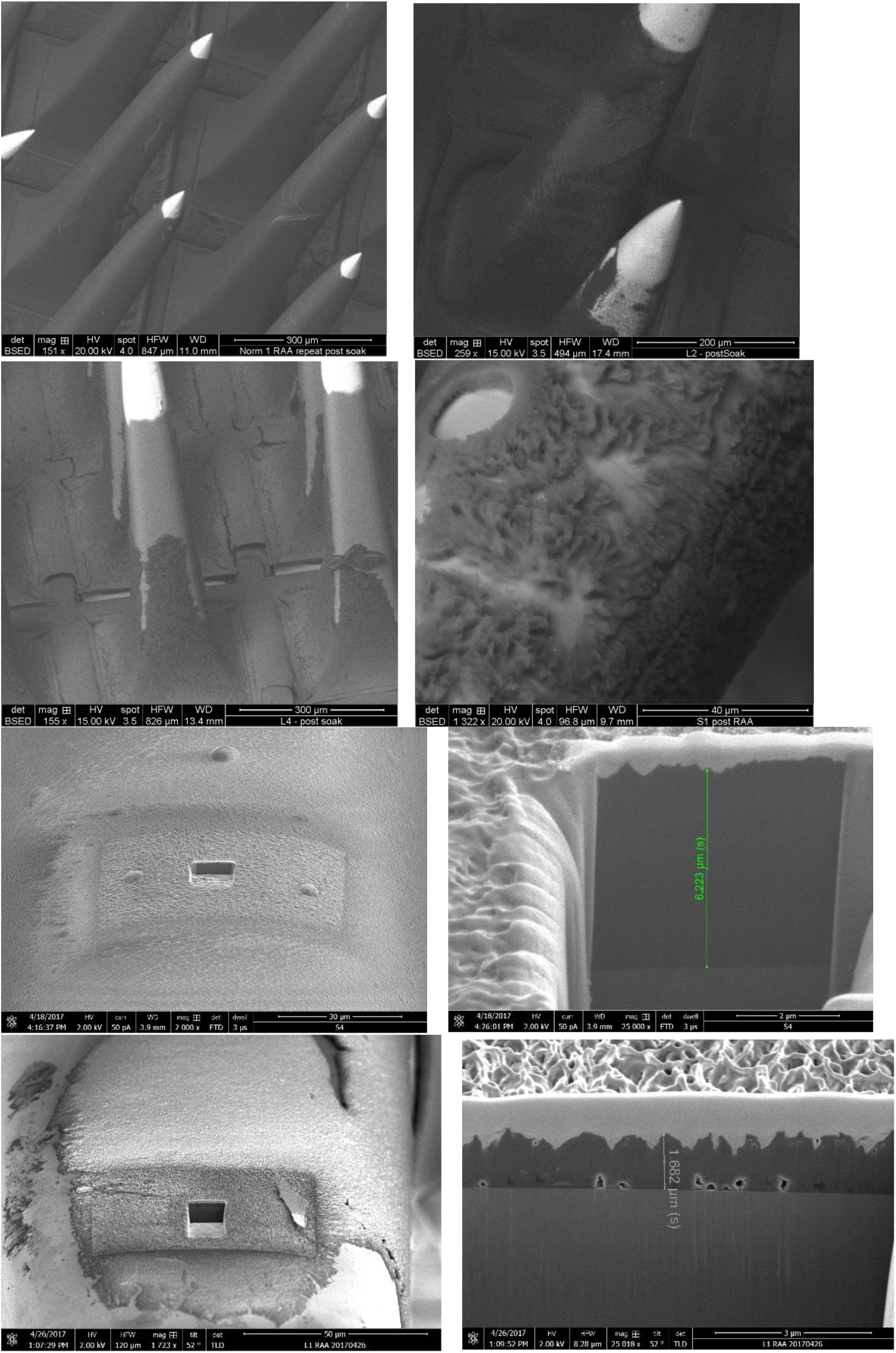
Select raw images used for Fig 5

**Fig. S5.**
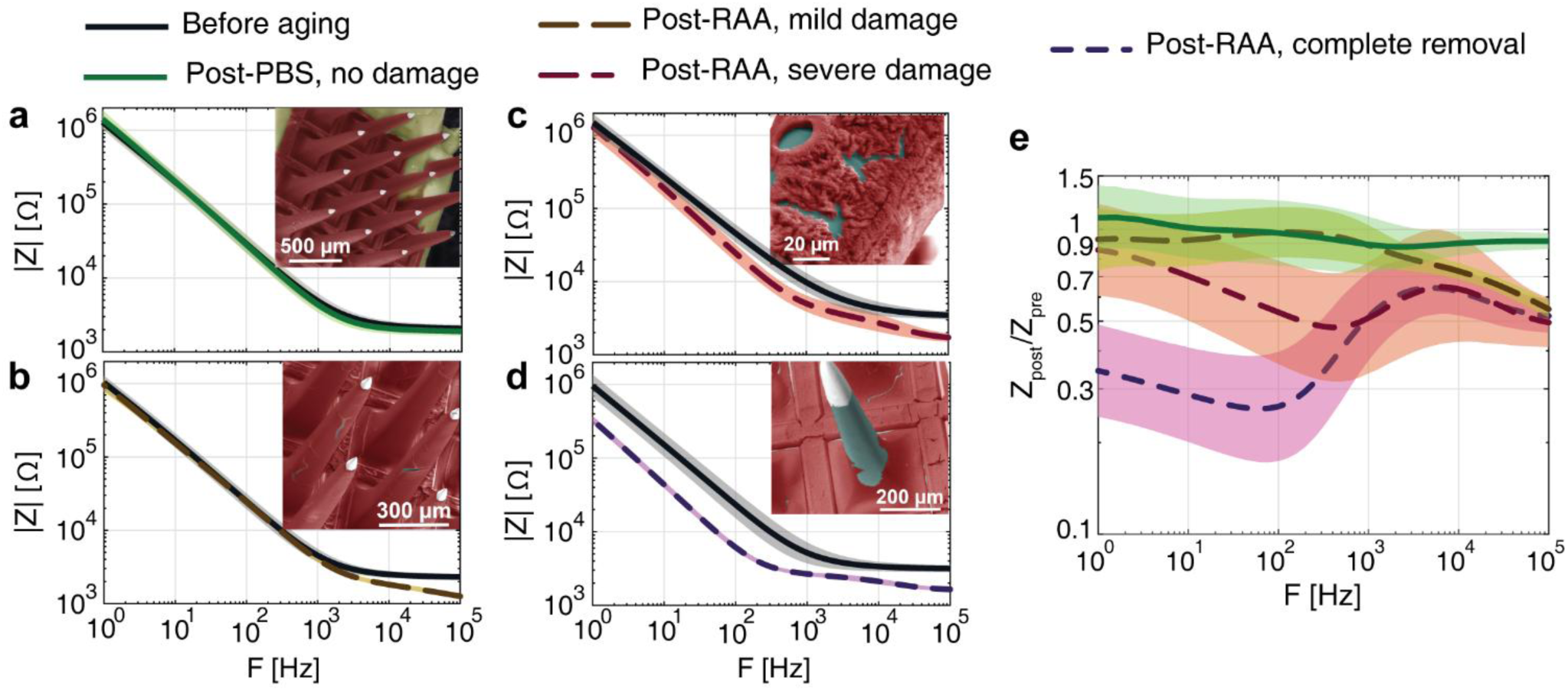
EIS results showing changes to EIS spectra commiserate with PPX-C damage. Insets are images of arrays from which spectra were taken. (a) No visible damage to PPX-C was seen for a control array aged at 87° C, corresponding with no meaningful change to impedance magnitude spectra. All remaining arrays were RAA-processed at 87 °C. (b) UEA with cracked parylene showed impedance reduction at frequencies >1000 Hz. (c) Degradation to parylene and more comprehensive shaft exposure from underneath encapsulation was accompanied by impedance reduction at 1000 Hz and below. (d) Full removal of PPX-C caused full-spectrum impedance reduction most pronounced at frequencies <1000 Hz. (e) Post-aging impedance normalized to initial values show how impedance change progressed across frequencies according to PPX-C damage mode. All plots are geometric averages of N=16 electrodes, shading indicates geometric standard deviation.

**Table S1.** EDS, XPS, FTIR sample numbers by device group.

| Device type | Processing condition | Sample number (N) |  |  |
| --- | --- | --- | --- | --- |
|  |  | EDS<sup>a</sup> | XPS<sup>b</sup> | FTIR<sup>b</sup> |
| Aged UEAs | *In vivo* | 12 | 2 | 4 |
|  | PBS, 67 °C | 14 | 1 | 4 |
|  | PBS, 87 °C | 15 | 3 | 4 |
|  | RAA, 67 °C | 18 | 1 | 4 |
|  | RAA, 87 °C | 20 | 3 | 4 |
| Aged test structures | PBS, 67 °C | 12 | 0 | 4 |
|  | PBS, 87 °C | 6 | 0 | 1 |
|  | RAA, 67 °C | 9 | 0 | 1 |
|  | RAA, 87 °C | 10 | 0 | 4 |
| UEA references | No deinsulation | 6 | 1 | 1 |
|  | Deinsulation | 10 | 1 | 2 |
| Planar references | Oxidized | 10 | 2 | 4 |
|  | O<sub>2</sub> etched | 3 | 1 | 1 |
|  | Untreated | 6 | 1 | 4 |

<sup>a</sup>N is sum of 2-3 measurements per device, save *in vivo* samples.<sup>b</sup>One measurement taken per device, save *in vivo* samples.

